# Impaired ERK MAPK activation in mature osteoblasts enhances bone formation via the mTOR pathway

**DOI:** 10.1101/2022.01.24.477465

**Authors:** Jung-Min Kim, Yeon-Suk Yang, Jaehyoung Hong, Sachin Chaugule, Hyonho Chun, Marjolein C. H. van der Meulen, Ren Xu, Matthew B. Greenblatt, Jae-Hyuck Shim

**Affiliations:** Department of Medicine, University of Massachusetts Chan Medical School, Worcester, MA, USA; Department of Mathematical Sciences, Korea Advanced Institute of Science and Technology, Daejeon, Republic of Korea; Meinig School of Biomedical Engineering and Sibley School of Mechanical & Aerospace Engineering, Cornell University, Ithaca, NY, USA; Research Division, Hospital for Special Surgery, New York, NY, USA; State Key Laboratory of Cellular Stress Biology, School of Medicine, Xiamen University, Xiamen, Fujian, China; Fujian Provincial Key Laboratory of Organ and Tissue Regeneration, School of Medicine, Xiamen University, Xiamen, Fujian, China; Department of Pathology and Laboratory Medicine, Weill Cornell Medical College, New York, NY, USA; Horae Gene Therapy Center, University of Massachusetts Chan Medical School, Worcester, MA, USA; Li Weibo Institute for Rare Diseases Research, University of Massachusetts Chan Medical School, Worcester, MA, USA

**Keywords:** MEK, ERK, mTOR, and osteoblast

## Abstract

Emerging evidence supports that osteogenic differentiation of skeletal stem cells (SSCs) is a key determinant of overall bone formation and bone mass. Despite extensive studies showing mitogen-activated protein kinase (MAPK) function in osteoblast differentiation, none of these studies properly show *in vivo* evidence of impacting post-lineage commitment and subsequent maturation. Here, we describe how the extracellular signal-regulated kinase (ERK) pathway in osteoblasts controls bone formation by suppressing the mechanistic target of rapamycin (mTOR) pathway. We also show that, while ERK inhibition blocks the differentiation of osteogenic precursors when initiated at an early stage, ERK inhibition surprisingly promotes the later stages of osteoblast differentiation. Accordingly, inhibition of the ERK pathway using a small compound inhibitor or conditional deletion of the MAP2Ks *Mek1* and *Mek2*, in mature osteoblasts and osteocytes (*Mek1/2^Dmp1^*), markedly increased bone formation due to augmented osteoblast differentiation. Mice with inducible deletion of the ERK pathway in mature osteoblasts (*Mek1/2^Ocn-Ert^*) also displayed similar phenotypes, demonstrating that this phenotype reflects continuous postnatal inhibition of late-stage osteoblast maturation. Mechanistically, ERK inhibition increases mitochondrial function and SGK1 phosphorylation via mTOR2 activation, which leads to osteoblast differentiation and production of angiogenic and osteogenic factors to promote bone formation. This phenotype was partly reversed by inhibiting mTOR. Our study uncovers a surprising dichotomy of ERK pathway functions in osteoblasts, whereby ERK activation promotes the early differentiation of osteoblast precursors, but inhibits the subsequent differentiation of committed osteoblasts via mTOR-mediated regulation of mitochondrial function and SGK1.

## Introduction

Signal transduction and transcription programs are coordinated to determine the commitment, proliferation, and differentiation of skeletal stem cells (SSCs) to osteoblasts. Once SSCs are committed to osteoprogenitors, they become pre-osteoblasts, proliferate, and differentiate into mature osteoblasts producing mineral and extracellular matrix proteins. Mature osteoblasts lastly terminally differentiate into osteocytes and embed themselves within the bone matrix (Salhotra, Shah et al., 2020). Proliferation is dominant at an early stage of osteogenesis, while proliferation rates decrease and extracellular mineralization increases during osteoblast differentiation (Infante & Rodriguez, 2018, Long, 2011). The balance between proliferation and differentiation during osteogenesis is tightly regulated by the sequential activation of signaling cascades, such as the mitogen-activated protein kinase (MAPK) and the mammalian/mechanistic target of rapamycin (mTOR) pathways.

The extracellular signal-regulated kinase (ERK) MAPK pathway transduces activation signals from extracellular cues, such as bone morphogenic proteins (BMPs) and wingless-related integration site (WNT) ligands, determining the specification of SSCs to osteoprogenitors (Ge, Xiao et al., 2007, Greenblatt, Shim et al., 2013, Kim, Yang et al., 2019, Matsushita, Chan et al., 2009). Conditional deletion of ERK1 and 2 MAPKs in the mesenchyme (*Erk1^-/-^Erk2^Prx1^*) causes severe limb deformity and bone defects due to impaired osteogenesis (Matsushita et al., 2009). Similarly, mice lacking the MAP2Ks in the ERK pathway, *Mek1* and *2*, in osteoprogenitors (*Mek1^Osx^Mek2^−/−^*) showed low bone mass and a cleidocranial dysplasia-like phenotype (CCD), similar to that seen in mice and humans with runt-related transcription factor 2 (*RUNX2)* haploinsufficiency (Ge et al., 2007, Ge, Xiao et al., 2009, Kim et al., 2019). Thus, ERK-mediated activation of RUNX2 is required for the commitment of SSCs to osteoprogenitors and the subsequent proliferation of osteoprogenitors at early differentiation stages. However, the role of ERK in the regulation of mature osteoblasts post-lineage commitment is unresolved (de la Croix Ndong, Stevens et al., 2015, Tang, Wu et al., 2021).

In addition to RUNX2, the ERK MAPK pathway controls the mTOR pathway, a signaling modulator of mitochondrial biogenesis that regulates cellular energy metabolism (Morita, Gravel et al., 2013). Cross-regulation between these pathways in various cancer cells has been also reported (Mendoza, Er et al., 2011). ERK inhibition by a MEK inhibitor activates the mTORC2- AKT signaling axis downstream of epidermal growth factor (EGF) (Yu, Liu et al., 2002), and a similar increase in AKT activation by ERK inhibition was reported in anaplastic lymphoma kinase (ALK)–addicted neuroblastoma (Umapathy, Guan et al., 2017). Thus, determining the function and the regulatory mechanism of ERK in osteoblast differentiation and bone formation is clinically significant as the ERK pathway is the most frequently mutated pathway in cancer and, accordingly, ERK pathway components, including rat sarcoma (RAS), rapidly accelerated fibrosarcoma (RAF), and MEK isoforms, are all well-established targets to treat a wide variety of ERK pathway-activated cancers (Hong, Fakih et al., 2020, Hyman, Puzanov et al., 2015, Robert, Karaszewska et al., 2015, Tiacci, De Carolis et al., 2021). Given that many of these patients are at an increased risk for fracture, it is important to weigh the risks and benefits when determining the role of the ERK pathway in the regulation of bone mass and the potential skeletal impact of ERK pathway-directed therapies.

Here, we establish dynamic roles for the ERK MAPK pathway during osteogenesis, with the ERK-mTOR signaling axis acting in osteoblasts to regulate mitochondria-dependent energy metabolism and SGK1-mediated production of pro-angiogenic and osteogenic factors. ERK inhibition in mature osteoblasts promotes bone formation via mTOR-mediated activation of mitochondria and SGK1. Accordingly, inhibition of the mTOR pathway using rapamycin attenuated bone accrual in *MEK1/2*-deficient mice. Our study uncovers a surprising dichotomy of ERK pathway functions in osteoblasts, with the ERK pathway promoting bone formation in osteoblast precursors but inhibiting bone formation in committed osteoblasts via mTOR pathway.

## Results

### A biphasic role for the ERK MAPK pathway in osteogenesis

To investigate the role of the ERK MPAK pathway in osteoporosis, the MEK inhibitor trametinib was treated to mice with estrogen deficiency-induced bone loss. 12-week-old mice were subjected to the ovariectomy (OVX) model of postmenopausal osteoporosis or sham surgery and were subsequently treated with trametinib via oral gavage for 8 weeks. While OVX surgery induced a significant bone loss in the femur of vehicle-treated mice, femoral bone loss was prevented by treatment with trametinib, as shown by a greater trabecular bone volume (Tb. BV) and trabecular thickness (Tb. Th) (Fig 1A). Remarkably, dynamic histomorphometry analysis demonstrated an increase in bone formation rate per bone surface (BFR/BS) and osteoblast surface on the trabecular bone area (Ob.S/BS) of trametinib-treated OVX mice (Fig 1B and C). Thus, ERK inhibition prevents estrogen deficiency-induced bone loss by promoting bone formation due to augmented osteoblast activity.

**Figure 1.**
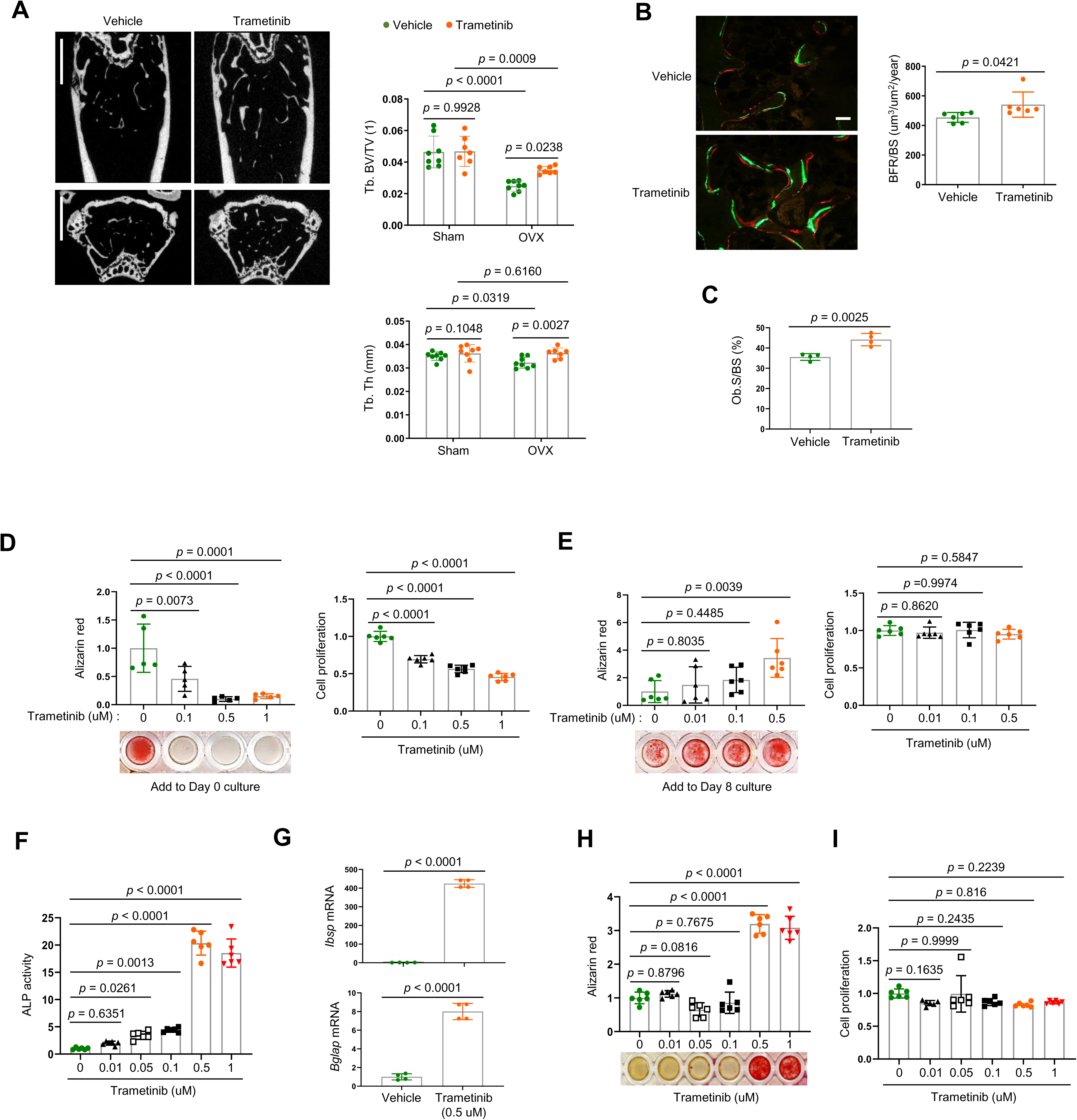
**The ERK MAPK pathway plays biphasic roles in osteogenesis** (A-C) Twenty-week-old female mice were treated with vehicle or trametinib after either Sham or OVX surgery. Representative microCT images of the femoral bone (A, left) and quantification of relative trabecular bone volume/total volume (Tb. BV) and thickness (Tb. Th, A, right) are displayed. Alternatively, dynamic histomorphometry analysis of the femoral bones was performed to assess *in vivo* osteoblast activity. Representative images of calcein/alizarin labeling (B, left) and quantification of bone formation rate (BFR)/bone surface (BS, B, right) and osteoblast surface (Ob.S)/bone surface (BS, C) are displayed. Scale bars, 1 mm (A), 50 μm (B). (D, E) Human bone marrow-derived mesenchymal stromal cells (BMSCs) were treated with different doses of trametinib at day 0 (D) or day 8 (E) of osteogenic culture. Mineralization and cell proliferation were assessed by alizarin red and alamar blue staining after 14 days of culture, respectively. (F-I) Mouse wildtype calvarial osteoblasts (COBs) were treated with different doses of trametinib at day 0 of osteogenic culture and 6 days later, ALP activity (F) and osteogenic gene expression by RT-PCR (G) were assessed. Mineralization by alizarin red staining (H) and cell proliferation (I) by alamar blue staining were assessed after 18 days of culture. Data are representative of three independent experiments (A [left], B [left], D-I) or pooled from two experiments (A [right], B [right], C). A two-tailed unpaired Student’s t-test for comparing two groups (A-C, G) or ordinary one-way ANOVA with Dunnett’s multiple comparisons test (D-F, H, I) (A-I; error bars, SD of biological replicates).

To gain insights into the mechanism underlying this finding, we examined the effects of ERK inhibition on the proliferation and differentiation of osteoblast precursors using human bone marrow-derived mesenchymal stromal cells (BMSCs). Expression of key osteogenic commitment factors, *RUNX2* and *SP7* (Osterix), was markedly upregulated 8 days after osteogenic culture as expected, establishing that the BMSCs at this time point represent committed osteoblast lineage cells (Fig EV1). ERK inhibition starting at day 8 of culture promoted osteoblast differentiation, whereas ERK inhibition from day 0 of culture conversely inhibited proliferation and osteogenic potential of BMSCs (Fig 1D and E). Similar findings were observed with committed osteoblasts isolated from mouse calvarium, which also showed increased osteogenic differentiation in the presence of trametinib. This was evident in ERK inhibitor induced increases in alkaline phosphatase activity (ALP), mineralization, and expression of osteogenic genes, including integrin-binding sialoprotein (*Ibsp*) and osteocalcin (*Bglap*) (Fig 1F-H). Notably, cell proliferation rates were largely unaffected (Fig 1I). These results suggest that while the ERK MAPK pathway plays a positive role in the early stages of osteogenesis, it acts as a negative regulator of the later stages of osteoblast differentiation. Thus, ERK inhibition in late-stage osteoblasts by trametinib treatment is likely to be responsible for the increased osteoblast activity observed in ERK inhibitor-treated OVX mice.

### Impaired ERK signaling in mature osteoblasts promotes bone formation

To confirm these findings using genetic approaches *in vivo*, *Mek1* and *Mek2,* which are upstream of ERK1/2, were conditionally deleted in mature osteoblasts and osteocytes by crossing mice bearing a floxed allele of *Mek1* (*Mek1^fl/fl^*) to the *Mek2*-null background (*Mek2*^-/-^) (Belanger, Roy et al., 2003, Bissonauth, Roy et al., 2006) (*Mek1^fl/fl^Mek2*^-/-^). These mice were then further crossed with *Dmp1* (Dentin matrix protein1)-*Cre* mice, which targets conditional gene deletion to mature osteoblasts and osteocytes (Lu, Xie et al., 2007). Deletion of *Mek1/2* was confirmed in RNA from the tibiae of WT and *Mek1^Dmp1^Mek2*^-/-^ (double knockout, *dKO^Dmp1^*) mice (Fig 2A), and resulted in a significant increase in femoral bone mass, as evidenced by greater trabecular bone volume (Tb. BV), thickness (Tb. Th), number (Tb. N), and cortical thickness (Ct. Th) (Fig 2B and C). Likewise, skull thickness and L4 vertebrae trabecular bone volume/tissue volume (Tb. BV/TV) were markedly increased (Fig 2D and E). These skeletal phenotypes require the deletion of both *Mek1* and *Mek2* alleles as deletion of *Mek1* (*Mek1^Dmp1^*) or *Mek2* (*Mek2*^-/-^) alone did not show an increase in bone mass compared to WT femurs (Fig 2C), suggesting a redundant role for *Mek1* and *Mek2* in osteoblasts. Consistent with the increased bone mass observed, a three-point bending test of *dKO^Dmp1^* mice revealed an increase in maximum bending moment and load at failure, demonstrating enhanced bone mechanical strength (Fig 2F). Thus, the bone formed by ERK pathway ablation in late-stage osteoblasts has the biomechanical properties of mature, physiologic bone and not that of fracture-prone pathologic bone. Finally, dynamic histomorphometry analysis demonstrated a significant increase in BFR/BS and mineral apposition rate (MAR) of *dKO^Dmp1^* femurs, indicating enhanced *in vivo* osteoblast activity in the absence of *Mek1/2* (Fig 2G). Therefore, deletion of *Mek1/2* in late-stage osteoblast lineage cells augmented their bone-forming activity and bone accrual and strength. Importantly, *in vivo* osteoclast activity and numbers were markedly elevated in *dKO^Dmp1^* mice, as shown by an increase in TRAP-positive osteoclasts, bone erosion surface, and serum levels of the bone resorption marker C-terminal telopeptide type I collagen (CTx-I) (Fig EV2A and B). These results suggest that elevated osteoblast activity caused by *Mek1/2*-deletion leads to increased osteoclast activity, thereby enhancing bone remodeling activity.

**Figure 2.**
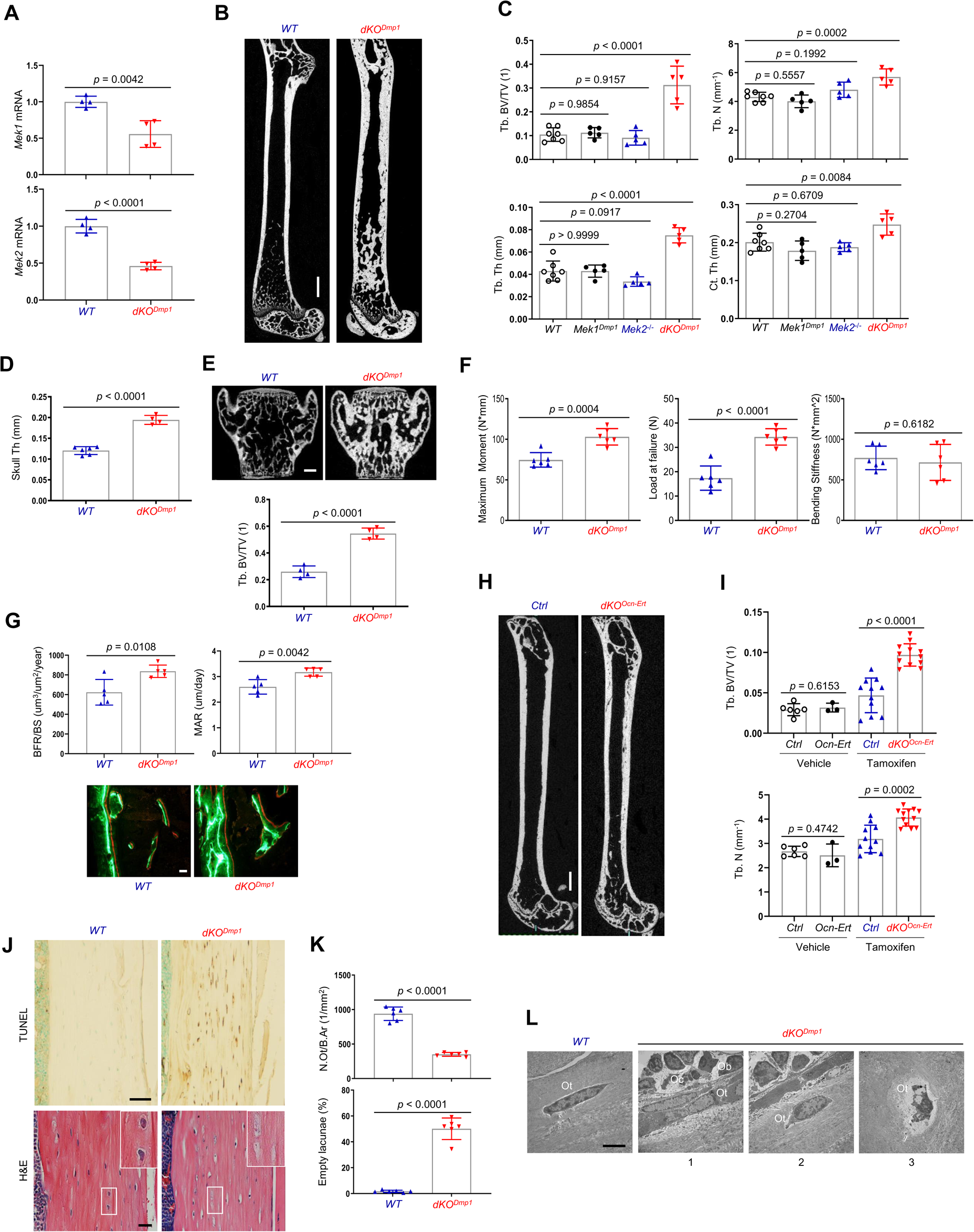
***Mek1/2* deletion in mature osteoblasts promotes bone formation in mice** (A) *Mek1* and *Mek2* mRNA levels in the tibial bones of *Mek1^fl/fl^Mek2^+/+^* (WT) and *Mek1^Dmp1^Mek2^-/-^* (*dKO^Dmp1^*) mice were measured by RT-PCR analysis. (B, C) MicroCT analysis showing femoral bone mass in eight-week-old *Mek1^fl/fl^Mek2^+/+^* (*WT*), *Mek1^Dmp1^Mek2^+/+^*(*Mek1^Dmp1^*), *Mek1^fl/fl^Mek2^-/-^* (*Mek2^-/-^*), and *Mek1^Dmp1^Mek2^-/-^* (*dKO^Dmp1^*) mice. 3D- reconstruction (B) and the relative quantification (C) are displayed. Trabecular bone volume/total volume (Tb. BV/TV), trabecular thickness (Tb. Th), trabecular number (Tb. N), and cortical thickness (Ct.Th). Scale bar, 1 mm (B). (D, E) MicroCT analysis of eight-week-old *WT* and *dKO^Dmp1^* skull (D) and vertebrae (E). Quantification of skull thickness (Th, D) and trabecular bone mass (E, bottom) and 3D- reconstruction of lumbar 4 (L4) vertebrae (E, top) are shown. Scale bar, 200 μm (E). (F) Biomechanical properties of eight-week-old *WT* and *dKO^Dmp1^* femurs, including maximum moment, load at failure, and bending stiffness were quantified using three-point bending test. (G) Dynamic histomorphometry analysis of eight-week-old *WT* and *dKO^Dmp1^* femurs. Quantification (top) and images of calcein/alizarin labeling (bottom) are displayed. BFR/BS, bone formation rate per bone surface; MAR, mineral apposition rate. Scale bar, 50 μm (bottom). (H, I) MicroCT analysis of femoral bone mass in seventeen-week-old *Mek1^fl/fl^Mek2^−/−^* and *Mek1^Ocn-Ert^Mek2^−/−^* mice treated with vehicle or tamoxifen; vehicle-treated *Mek1^fl/fl^Mek2^−/−^* (*Ctrl*) and *Mek1^Ocn-Ert^Mek2^−/−^* (*Ocn-Ert*) mice, tamoxifen-treated *Mek1^fl/fl^Mek2^-/-^* (*Ctrl*) and *Mek1^Ocn- Ert^Mek2^−/−^*(*dKO^Ocn-Ert^*) mice. 3D-reconstruction (H) and the relative quantification (I) are displayed. Scale bar, 1 mm (H). (J) TUNEL (top) and H&E (bottom)-stained longitudinal sections of eight-week-old *WT* and *dKO^Dmp1^* femurs. Scale bars, 50 μm (top) and 20 μm (bottom). (K) Numbers of osteocytes/bone area (N.Ot/B.Ar) and empty lacunae in the cortical bones of eight- week-old *WT* and *dKO^Dmp1^* femurs. (L) Transmission electron microscopy (TEM) images of osteocytes in the cortical bones of eight- week-old *WT* and *dKO^Dmp1^* femurs. 1, Ot in osteoids; 2, Ot in mineralized bone matrix; 3, apoptotic Ot in bone matrix. Ot, osteocyte; Oc, osteoclast; Ob, osteoblast. Scale bar, 5 μm. Data are representative of three independent experiments (A, B, E [top], G [bottom], H, J, L) or pooled from two experiments (C, D, E [bottom], F, G [top], I, K). A two-tailed unpaired Student’s t-test for comparing two groups (A, D-G, I, K) or ordinary one-way ANOVA with Dunnett’s multiple comparisons test (C) (A, C-G, I, K; error bars represent the SD of biological replicates).

To confirm that deletion of *Mek1/2* in mature osteoblasts is responsible for the development of these phenotypes, we generated inducible, osteoblast-specific *Mek1/2*-knockout mice by crossing *Mek1^fl/fl^Mek2*^-/-^ mice with osteocalcin-*CreERT* mice expressing a tamoxifen-induced *Cre* recombinase in mature osteoblasts (Maes, Kobayashi et al., 2010) (*dKO^Ocn-Ert^*). Treatment of *dKO^Ocn-Ert^* mice with tamoxifen significantly increased trabecular bone mass of the femur (Fig 2H and I), demonstrating that the inhibitory role of the ERK MAPK pathway is specific to mature osteoblasts and reflects the continuous function of ERK in these cells and is not secondary to a developmental abnormality.

Given previous studies showing the importance of the ERK MAPK pathway in osteocyte survival (Plotkin, Mathov et al., 2005, Ru & Wang, 2020), terminal deoxynucleotidyl transferase dUTP nick end labeling (TUNEL) staining was performed in *dKO^Dmp1^* femurs to assess osteocyte cell death (Fig 2J-top). Osteocyte apoptosis was substantially increased in the cortical bone of *dKO^Dmp1^* mice relative to control mice, which corresponded to decreased osteocyte numbers and increased empty lacunae (a cavity within the bone) (Fig 2J-bottom and K). Likewise, transmission electron microscopy (TEM) analysis of *dKO^Dmp1^* femurs demonstrated that terminally differentiated osteocytes embedded in the bone matrix undergo apoptosis [Fig 2L (3)] while osteocytes at early [Fig 2L (1)] and intermediate [Fig 2L (2)] differentiation stages were grossly normal in the absence of *Mek1/2* (Fig 2L). This is consistent with RT-PCR analysis showing that mRNA levels of early osteocyte markers, including dentin matrix protein1 (*Dmp1*) and phosphate regulating endopeptidase homolog X-linked (*Phex*) were comparable between control and *dKO^Dmp1^* tibias. However, expression of the late osteocyte marker, sclerostin (*Sost*), was significantly decreased (Fig EV2C). These results suggest that ERK activation is required for the survival of late/fully mature osteocytes, but not the transition of mature osteoblasts to early osteocytes. Taken together, these results demonstrate that ERK inhibition in mature osteoblasts promotes bone formation, while impaired ERK signaling in late osteocytes results in cell death.

### ERK inhibition increases osteogenesis and production of angiogenic and osteogenic factors

As seen in *dKO^Dmp1^* mice, CRE-induced deletion of *Mek1* and *Mek2* MAP2Ks in committed osteoblasts increased ALP activity, mineralization, and osteogenic gene expression. Deletion of *Mek1* or *Mek2* alone did not impact osteogenic differentiation (Figs 3A-C and EV3A). While proliferation rates of control osteoblasts were markedly increased under osteogenic conditions, little to no increase was observed in the absence of *Mek1/2* (Fig 3D), suggesting that ERK inhibition may control a shift towards differentiation at the expense of proliferation. To gain insights into the mechanisms by which ERK inhibition promotes osteoblast differentiation, we performed whole transcriptome analysis of *dKO* osteoblasts six days after cell growth (GM) or osteogenic (OIM) culture conditions. Gene ontology analysis demonstrated that, under cell growth conditions, upregulated genes were highly enriched in the pathways involved in ossification, skeletal system morphogenesis, and connective tissue development (Fig 3E), consistent with an increase in ALP activity and expression of osteogenic genes in undifferentiated *dKO* osteoblasts (Fig EV3B and C). These results suggest that ERK inhibition induces spontaneous osteogenic differentiation by upregulating transcription of genes favorable to osteogenic differentiation. Under osteogenic conditions, gene ontology analysis of differentiated *dKO* osteoblasts revealed upregulation of genes associated with extracellular matrix organization, while genes related to cell proliferation, including nuclear division and cell cycle, were downregulated (Figs 3F and EV3D-G). These results suggest that impaired ERK signaling in later stage committed osteoblasts upregulates transcriptional programs associated with osteogenic differentiation while suppressing the induction of genes associated with cell proliferation.

**Figure 3.**
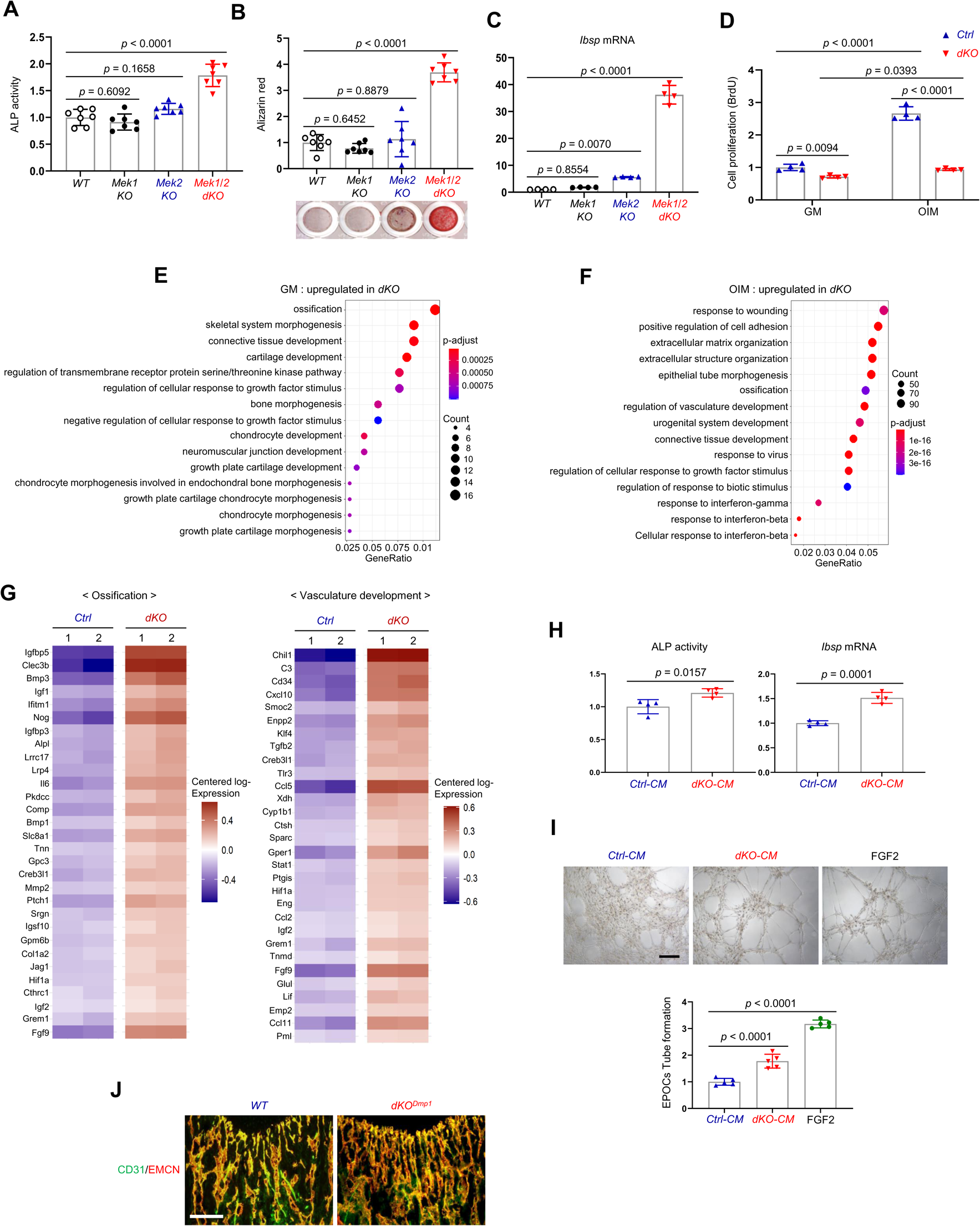
**Effects of *Mek1/2* deletion on osteoblast differentiation and production of angiogenic and osteogenic factors** (A-C) Mouse *Mek1^fl/fl^Mek2^+/+^* and *Mek1^fl/fl^Mek2^-/-^* COBs were infected with control vector or CRE recombinase-expressing lentiviruses; *Mek1^fl/fl^Mek2^+/+^* COBs with control (WT) or CRE (*Mek1 KO)*, *Mek1 ^fl/fl^Mek2^-/-^* COBs with control (*Mek2 KO)* or CRE *(Mek1/2 dKO)*. Puromycin-selected COBs were cultured under osteogenic conditions and ALP activity (A) and osteogenic gene expression (C) were determined at day 6 and mineralization (B) was analyzed at day 18 of culture. (D) *Mek2 KO* (*Ctrl*) and *Mek1/2 dKO (dKO)* COBs were cultured with control growth medium (GM) or osteogenic induction medium (OIM) and cell proliferation was analyzed by BrdU incorporation at day 6 of the culture. (E, F) Transcriptome analysis of *Ctrl* and *dKO* COBs 6 days after GM or OIM culture. Biological process output of gene ontology analysis was performed in both GM and OIM group for upregulated genes in *dKO* relative to *Ctrl* COBs. The color indicates adjusted P-value as estimated by the Benjamini–Hochberg method with the threshold of significance *p* = 0.05 and *q* = 0.005. (G) Heatmaps for ossification- and vasculature-associated gene expression. The top 30 upregulated genes in *dKO* COBs relative to *Ctrl* COBs are displayed as each row and column represent gene symbol and sample, respectively. The log10 expression (read count) was centered across samples and red and purple denote upregulated and downregulated, respectively. (H) Conditioned medium (CM) from *Ctrl* and *dKO* COBs were collected at day 6 under osteogenic culture and mouse wildtype BMSCs were cultured under osteogenic condition in the presence of CM of *Ctrl* COBs and *dKO* COBs and ALP activity (left) and *Ibsp* mRNA level (right) were assessed at day 6. (I) Capillary tube formation of mouse endothelial cells (EPOCs) was performed in the presence of CM for 5 hours. Representative images (top) and quantification for the number of branches are displayed (bottom). FGF2 was used as a positive control. Scale bar, 200 μm. (J) Immunofluorescence for CD31 (green) and endomucin (EMCN, red) in the epiphyseal area of eight-week-old *WT* and *dKO^Dmp1^* femurs. Scale bar, 100 μm. Data are representative of three independent experiments (A-D, H-J). For transcriptome analysis, biological duplicates were analyzed (E-G). Ordinary one-way ANOVA with Dunnett’s multiple comparisons test (A-C, I) or a two-tailed unpaired Student’s t-test for comparing two groups (D, H) (A-D, H, I; error bars, SD of biological replicates).

Whole transcriptome analysis of *dKO* osteoblasts also revealed significant upregulation of secreted cytokines and/or growth factors involved in ossification and angiogenesis/vasculature development (Fig 3F), such as fibroblast growth factor 9 (*Fgf9*) (Behr, Leucht et al., 2010), ectonucleotide pyrophosphatase/phosphodiesterase 2 (*Enpp2*) (Cholia, Nayyar et al., 2015), transforming growth factor-beta 2 (*Tgfb2*) (Wu, Chen et al., 2016), insulin-like growth factors 1, 2 (*Igf1, 2*) (Yakar, Werner et al., 2018), bone morphogenic proteins 1, 3 (*Bmp1, 3*) (Wu et al., 2016), C-C Motif Chemokine Ligand 5 (*Ccl5*) (Suffee, Hlawaty et al., 2012), and *Ccl11* (Salcedo, Young et al., 2001) (Fig 3G). Accordingly, conditioned medium (CM), collected from *dKO* osteoblasts, markedly increased osteogenic potentials of wildtype BMSCs (Fig 3H) and capillary tube formation of bone marrow-derived endothelial progenitor outgrowth cells (EPOCs, Fig 3I). Likewise, an immunofluorescence analysis of *dKO^Dmp1^* femurs showed elevated levels of CD31- and endomucin-positive skeletal vasculature supporting osteoblast development (Xu, Yallowitz et al., 2018) (Fig 3J). These results suggest that pro-angiogenic and/or -osteogenic factors secreted from *dKO* osteoblasts enhance angiogenesis and osteogenesis, promoting bone formation. Taken together, *Mek1/2* deficiency does not only increase osteoblast differentiation by upregulating osteogenic transcriptional programs, but also improves bone-forming environment by producing pro-angiogenic and/or -osteogenic factors.

### ERK inhibition enhances mitochondrial energy metabolism in osteoblasts

It has been well-established that the ERK MAPK pathway controls cellular metabolism in cancer cells and highly proliferating stem cells (Lee, Guntur et al., 2017, Papa, Choy et al., 2019). Mature osteoblasts produce collagen for extracellular matrix formation using adenosine triphosphate (ATP) as a major energy resource (Lee et al., 2017). Moreover, glucose and glutamine, used for mitochondrial ATP production, are important for cell proliferation at early stages of osteoblast differentiation and matrix mineralization at later differentiation stages (Karner, Esen et al., 2015, Wei, Shimazu et al., 2015). Thus, we hypothesized that ERK-mediated regulation of glucose/glutamine metabolism may be involved in the determination of osteogenic potential and differentiation. Remarkably, the differentiation of *dKO* osteoblasts was substantially reduced by glutaminase inhibition but only modestly impacted by glucose deprivation (Fig 4A and B), demonstrating the importance of glutamine metabolism in ERK-mediated regulation of osteogenesis. Specifically, the responsiveness of *dKO* osteoblasts to the glutaminase inhibitor (bis- 2-[5-phenylacetamido-1,3,4-thiadiazol-2-yl]ethyl sulfide or BPTES) was greater than that of control osteoblasts, as shown by a significant reduction in ALP activity, cell proliferation, and osteogenic gene expression in the presence of BPTES (Fig 4B and C). Likewise, these cells showed elevated levels of glutaminase (*Gls*), a key enzyme that produces glutamate from glutamine (Fig 4D), and increased glutamate production and release (Fig 4E), suggesting enhanced glutamine metabolism in *dKO* osteoblasts. Of note, in contrast to previous studies showing critical roles of glucose metabolism (Wei et al., 2015) and the ERK MAPK pathway (Ge et al., 2007, Greenblatt et al., 2013, Kim et al., 2019, Matsushita et al., 2009) in controlling post-translational modifications of RUNX2 during early osteogenesis, RUNX2 expression and transcriptional activity were both intact in the absence of *Mek1/2* (Fig EV4), accompanied with little to no effect of glucose depravation on osteoblast maturation (Fig 4A). These results suggest that the ERK MAPK pathway is not required for glucose metabolism and RUNX2 post-translational modifications during the later stages of osteoblast differentiation. This finding provides a clear mechanistic distinction between the role of the ERK pathway in early osteogenic commitment which relates to regulation of RUNX2 activity, from the role of the ERK pathway in later stage osteoblasts which, by contrast, includes regulation of cellular energy metabolism.

**Figure 4.**
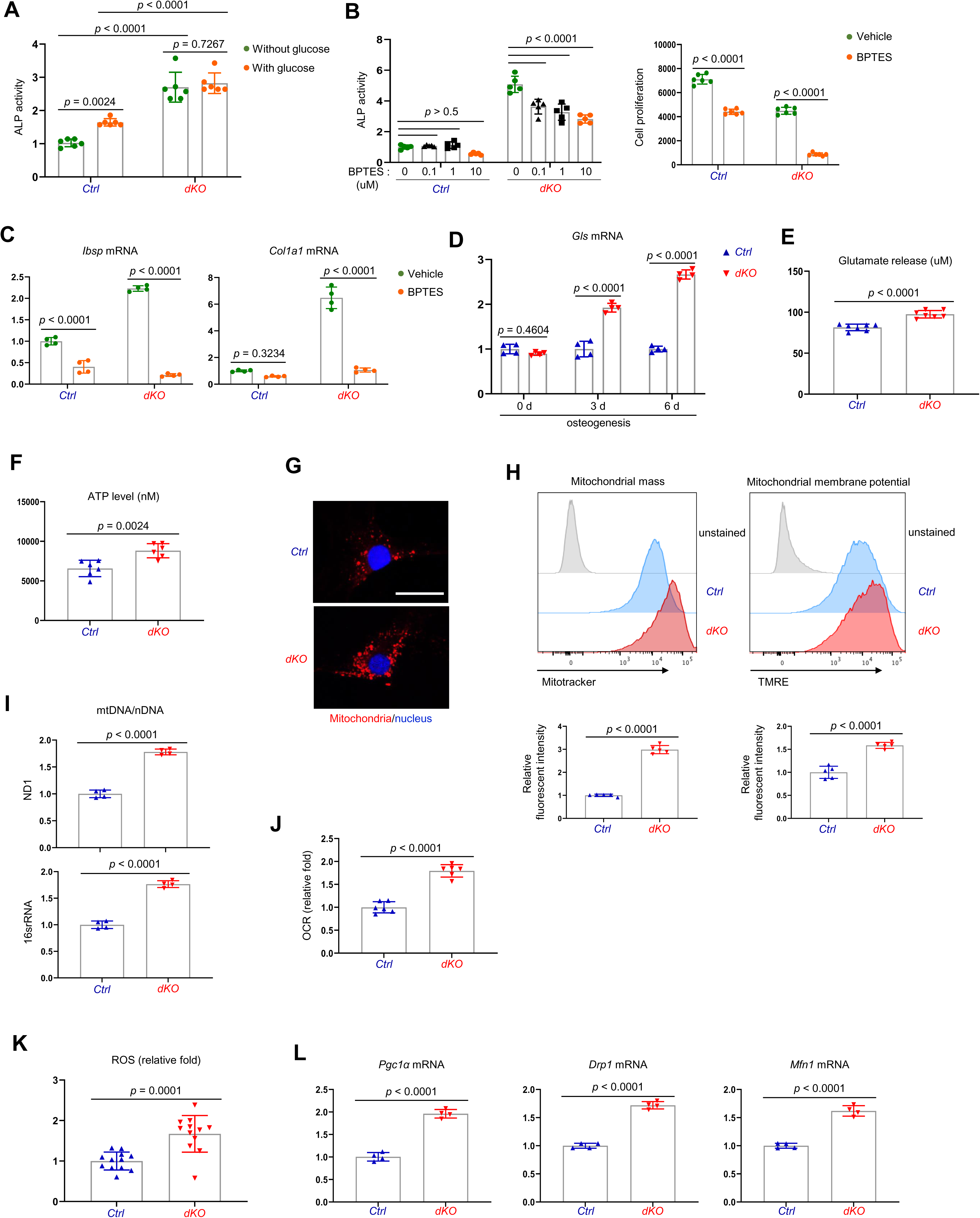
***Mek1/2* deletion enhances mitochondria-mediated energy metabolism in osteoblasts** (A) *Ctrl* and *dKO* COBs were cultured under osteogenic conditions in the presence or absence of glucose and 6 days later, ALP activity was assessed. (B, C) *Ctrl* and *dKO* COBs were treated with different doses of BPTES under osteogenic conditions and 6 days later, ALP activity and cell proliferation (B) and osteogenic gene expression (C) were assessed. (D, E) mRNA levels of *Gls* in *Ctrl* and *dKO* COBs were measured by RT-PCR analysis at different time points of osteogenic culture (D) and levels of glutamate in the supernatant were measured for extracellular glutamate release at day 6 culture of osteogenesis (E). (F-I) Intracellular ATP levels (F), mitochondrial numbers (G), and the ratio of mitochondrial DNA (mtDNA, mt-ND1 or mt-16sRNA) to nuclear DNA (nDNA, *Hk2*) (I) were assessed in *Ctrl* and *dKO* COBs after 6 days of culture. Alternatively, flow cytometry was used to measure mitochondrial numbers and membrane potential of *Ctrl* and *dKO* COBs treated with Mitotracker or TMRE at day 4 of osteogenic culture, respectively (H). Numbers indicate median fluorescence intensity (H, bottom). Scale bar, 75 μm (G). (J) Oxygen consumption rate (OCR) in *Ctrl* and *dKO* COBs 6 days after osteogenic culture. (K) Intracellular ROS levels in *Ctrl* and *dKO* COBs were assessed 6 days after osteogenic culture. (L) mRNA levels of mitochondria-related genes in *Ctrl* and *dKO* COBs were assessed by RT-PCR analysis. Data are representative of three independent experiments. A two-tailed unpaired Student’s t-test for comparing two groups (A, B [right], C-F, H-L) or ordinary one-way ANOVA with Dunnett’s multiple comparisons test (B [left]) (A-F, H-L; error bars, SD of biological replicates).

Given the importance of glutamine metabolism in mitochondria-mediated energy production, we next examined the intracellular ATP content in *dKO* osteoblasts and found increased ATP levels (Fig 4F). Additionally, mitochondrial numbers and membrane potentials and a ratio of mitochondrial DNA (mtDNA) levels to nuclear DNA (nDNA) levels in *dKO* or trametinib-treated osteoblasts were all markedly increased compared to control or vehicle-treated osteoblasts (Figs 4G-I and EV5). This corresponds to enhanced oxygen consumption rate (OCR), an indicator of cellular respiration, in *dKO* osteoblasts (Fig 4J), demonstrating that ERK inhibition promotes mitochondrial function and metabolic activity in committed osteoblasts. Finally, levels of reactive oxidative species (ROS) (Fig 4K) and mRNA levels of peroxisome proliferator-activated receptor gamma coactivator 1-alpha (*Pgc1α*), dynamin-related protein 1 (*Drp1*), and mitofusin 1 (*Mfn1*) (Fig 4L), genes that are important for mitochondrial biogenesis, fission, and fusion, were also markedly elevated in *dKO* osteoblasts. Taken together, these results suggest that ERK inhibition increases osteoblast differentiation via augmented glutamine metabolism and mitochondrial function.

### ERK inhibition promotes bone formation via the mTOR pathway in osteoblasts

As a metabolic hub in response to nutrients and growth factors, the mTOR pathway plays a crucial role in mitochondria-mediated cell metabolism during skeletal development and homeostasis (Chen & Long, 2018). Given that ERK functions upstream of the mTOR pathway in regulating mitochondria-mediated energy metabolism (Morita et al., 2013), we tested the ability of mTOR inhibition to reverse the enhanced osteogenic differentiation seen after *Mek1/2* deletion (Fig 5A). Rapamycin was used to inhibit two main mTOR signaling complexes, mTORC1 and mTORC2 (Lamming, Ye et al., 2012) in *dKO* osteoblasts, demonstrating that rapamycin treatment attenuated ALP activity (Fig 5B) and expression of osteogenic genes (Fig 5C) in the absence of *Mek1/2.* Of note, in contrast to many contexts where rapamycin solely inhibits the mTORC1 pathway, including p70S6 kinase (p70S6K), rapamycin treatment was effective at inhibiting the phosphorylation of the mTORC2 downstream molecules, protein kinase B (AKT) and serum/glucocorticoid regulated kinase 1 (SGK1), in *dKO* osteoblasts (Fig EV6). These results suggest that rapamycin treatment decreases the differentiation of *dKO* osteoblasts by inhibiting both mTORC1 and mTOCR2 pathways. To test this hypothesis *in vivo*, 2-week-old *Mek1^Dmp1^Mek2*^-/-^ (*dKO^Dmp1^*) mice were treated with rapamycin for 6 weeks and femoral bone mass was assessed in 8-week-old mice using microCT. While little effect of rapamycin on femoral bone mass was seen in control mice, trabecular bone mass and cortical bone thickness were both decreased in *dKO^Dmp1^* mice when treated with rapamycin relative to vehicle, as shown by a significant reduction in trabecular bone volume (Tb. BV), number (Tb. N), thickness (Tb. Th), and cortical thickness (Ct.Th) (Fig 5D and E). These results suggest, as observed in *dKO* osteoblasts, mTOR inhibition can reverse the enhanced bone-forming activity by *Mek1/2*-deletion in mice. A histologic analysis of rapamycin-treated *dKO^Dmp1^* femurs revealed that mTOR inhibition was also effective in preventing the osteocyte apoptosis occurring in the absence of *Mek1/2* (Fig 5F). Additionally, osteoclast surface (Oc.S) in the trabecular bone was slightly decreased in these femurs, suggesting that rapamycin treatment may directly suppress osteoclast differentiation and activity (Glantschnig, Fisher et al., 2003, Kneissel, Luong-Nguyen et al., 2004) (Fig 5G). Taken together, impaired ERK signaling is likely to promote osteogenic differentiation and bone formation via activation of the mTOR pathway, whose inhibition can reverse *dKO^Dmp1^* phenotypes *in vitro* and *in vivo*.

**Figure 5.**
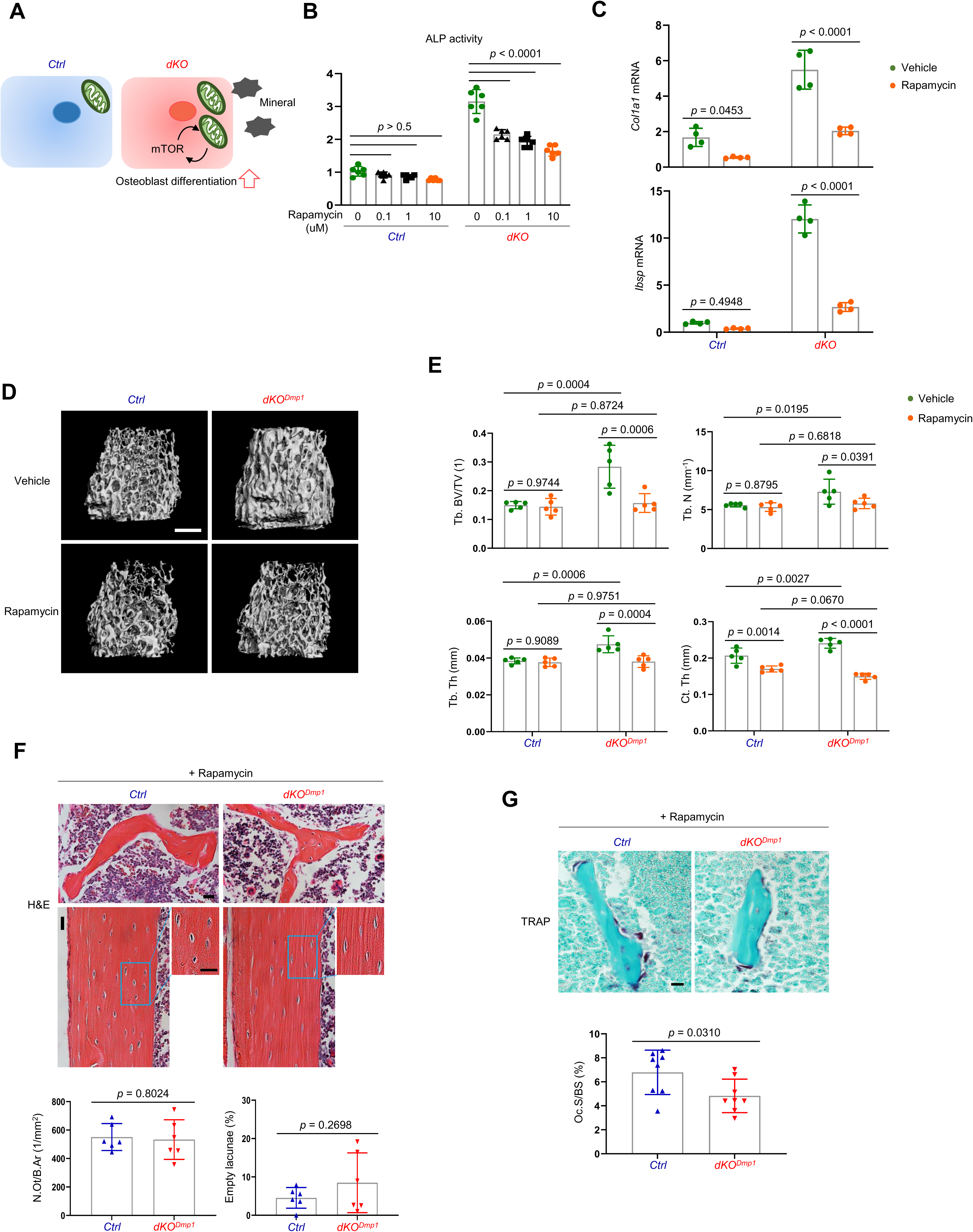
**Rapamycin treatment reverses skeletal phenotypes of *Mek1/2*-deficient mice** (A) Diagram showing the mechanisms by which ERK inhibition enhances osteoblast differentiation due to augmented mTOR-mediated mitochondrial function. (B, C) *Ctrl* and *dKO* COBs were treated with different doses of rapamycin under osteogenic conditions and 6 days later, ALP activity (B) and osteogenic gene expression (C) were determined. (D, E) MicroCT analysis showing femoral bone mass in eight-week-old mice treated with vehicle or rapamycin. 3D-reconstruction (D) and the relative quantification (E) are displayed. Scale bar, 500 μm (D). (F) H&E-stained longitudinal sections of eight-week-old *WT* and *dKO^Dmp1^* femurs treated with rapamycin. Representative images (top) and numbers of osteocytes/bone area (N.Ot/B.Ar) and empty lacunae (bottom) are displayed. Scale bar, 20 μm (top). (G) TRAP-stained longitudinal sections of eight-week-old *WT* and *dKO^Dmp1^* femurs treated with rapamycin. Representative images (top) and osteoclast surface/bone surface (Oc.S/BS) (bottom) are displayed. Scale bar, 20 μm (top). Data are representative of two or three independent experiments (B-D, F[top], G [top]) or pooled from two experiments (E, F [bottom], G [bottom]). Ordinary one-way ANOVA with Dunnett’s multiple comparisons test (B) or a two-tailed unpaired Student’s t-test for comparing two groups (C, E-G) (B, C, E-G; error bars, SD of biological replicates).

### ERK inhibition promotes osteoblast differentiation via mTORC2/SGK1 activation

To understand how ERK controls mTOR activation in osteoblasts, we examined the effects of *Mek1/2* deficiency on mTORC1 and mTORC2 signaling. While mTORC1 primarily promotes cell growth via activation of p70S6K and inactivation of eIF4E binding protein 1 (4EBP1), mTORC2 regulates cell survival and proliferation via AKT phosphorylation as well as cellular metabolism via activation of SGK1 and N-Myc downstream regulated 1 (NDRG1) (Saxton & Sabatini, 2017). In comparison to control osteoblasts, *dKO* osteoblasts showed a significant increase in phosphorylation levels of mTORC2 downstream molecules, including AKT, SGK1, and NDRG1, whereas little to no alteration in the phosphorylation levels of the mTORC1 downstream molecules p70S6K and 4EBP1 were detected in the absence of *Mek1/2* (Fig 6A). These results demonstrate that ERK inhibition in osteoblasts upregulates mTORC2- but not mTORC1-mediated signal transduction.

**Figure 6.**
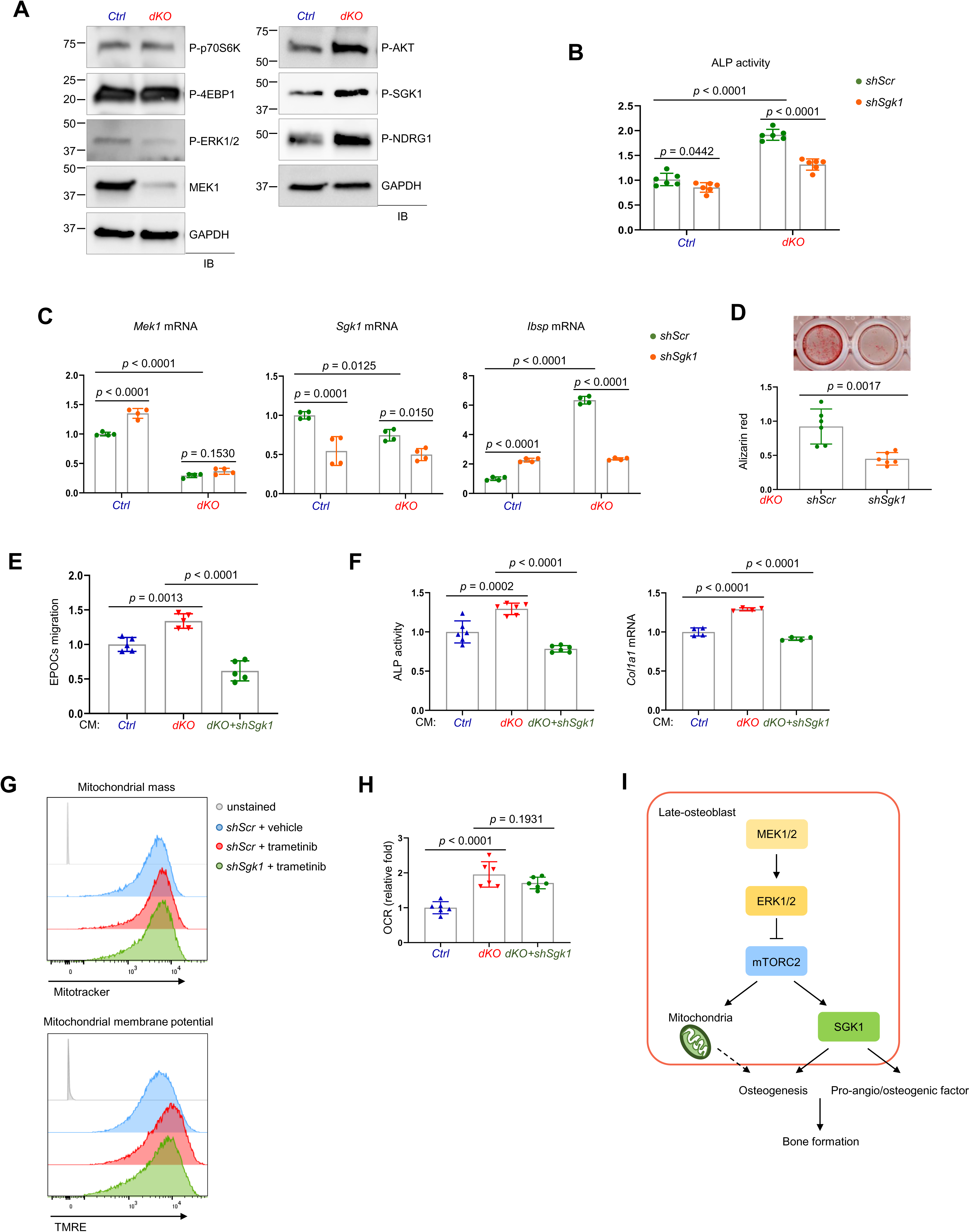
**mTORC2/SGK1 activation is important for *Mek1/2* deficiency-induced osteoblast differentiation** (A) Immunoblot analysis showing phosphorylation levels of mTOR signaling components in *Ctrl* and *dKO* COBs. GAPDH was used as a loading control. (B-D) *Ctrl* and *dKO* COBs expressing *shScr* or *shSgk1* shRNAs were cultured under osteogenic conditions. ALP activity (B), osteogenic markers expression (C), and mineralization (D) were assessed after 6 or 18 days of culture, respectively. (E) Conditioned medium (CM) collected from *Ctrl* and *dKO* COBs expressing *shScr* or *shSgk1* shRNAs 6 days after osteogenic culture was added to transwell migration of mouse endothelial cells (EPOCs) and migrated cells were assessed 12 hours after incubation. (F) Mouse BMSCs were cultured under osteogenic condition in the presence of CM, and ALP activity and *Cola1* mRNA level were analyzed at day 6 of the culture. (G) Vehicle- or trametinib-treated COBs expressing *shScr* or *shSgk1* shRNAs were treated with Mitotracker or TMRE 4 days after osteogenic culture and mitochondrial mass and membrane potential were assessed using flow cytometry, respectively. (H) OCR in *Ctrl* and *dKO* COBs expressing *shScr* or *shSgk1* shRNAs after 6 days of culture. (I) Diagram showing the molecular actions of ERK on bone formation. Data are representative of two or three independent experiments (A-H). A two-tailed unpaired Student’s t-test for comparing two groups (B-D) or ordinary one-way ANOVA with Dunnett’s multiple comparisons test (E, F, H) (B-F, H; error bars, SD of biological replicates).

In a previous study, SGK1, that functions downstream of mTORC2, played an important role in osteogenic trans-differentiation and calcification of vascular muscle cells (Voelkl, Luong et al., 2018). Since the genes associated with the ossification pathway in *dKO* osteoblasts were highly upregulated (Fig 3E and F), we hypothesized that ERK inhibition may upregulate osteoblast differentiation by activating the mTORC2-SGK1 signaling axis. Similar to rapamycin-mediated mTOR inhibition (Fig 5B and C), *Sgk1* knockdown markedly attenuated ALP activity (Fig 6B), expression of osteogenic genes (Fig 6C), and mineralization (Fig 6D) in *dKO* osteoblasts, not in control osteoblasts, suggesting that SGK1 functions downstream of ERK during osteoblast differentiation. Additionally, the ability of conditioned medium collected from *dKO* osteoblasts to promote angiogenesis of EPOCs and osteogenesis of wildtype BMSCs was ablated by *Sgk1* deficiency, as shown by a significant decrease in migration of EPOCs (Fig 6E), and ALP activity and *Col1a1* mRNA levels of BMSCs (Fig 6F). Notably, little to no alteration of mitochondrial numbers and membrane potentials (Fig 6G) and OCR (Fig 6H) was observed in trametinib-treated *Sgk1*-deficient osteoblasts or *Sgk1*-deficient *dKO* osteoblasts, suggesting that SGK1 is dispensable for ERK-mediated mitochondrial function. Taken together, the ERK-mTORC2 signaling axis mediates osteogenesis and production of pro-angio/osteogenic factors via SGK1 activation, while controlling osteogenesis via mitochondria-dependent energy metabolism (Fig 6I).

## Discussion

This study identifies a biphasic role for the ERK MAPK pathway in osteogenesis. While ERK activation is crucial for cell proliferation and osteogenic potential at the early stages of osteogenic differentiation, ERK-mediated regulation of the mTOR pathway determines osteogenic differentiation in later stages. In addition, ERK activation dampens the secretion of pro-angiogenic and osteogenic factors required to create an optimal microenvironment for bone formation. Mechanistically, impaired ERK signaling in committed osteoblasts enhances osteoblast differentiation via an increase in mitochondria-mediated metabolic demands, activation of mTORC2-SGK1 signaling axis, thereby promoting bone formation (Fig 6I). This corresponds to previous reports showing that mTORC2 activation is required for mitochondrial function and homeostasis (Wang, Shao et al., 2020) and that both glycolysis and oxidative phosphorylation by mitochondria are required to meet the energy demands needed for bone formation by osteoblasts (Lee et al., 2017). However, despite previous studies showing the importance of SGK1 in osteogenic trans-differentiation of vascular smooth muscle cells (Voelkl et al., 2018) and energy metabolism in different cellular types and functions (Mason, Cockfield et al., 2021), our findings suggest that while ERK-mTORC2-SGK1 signaling axis is required for osteogenesis and production of pro-angiogenic and -osteogenic factors, it is dispensable for mitochondrial function. Thus, our study identified two distinct pathways downstream of the ERK-mTORC2 signaling axis controlling bone formation, mitochondria-mediated osteogenesis and SGK1-mediated osteogenesis and production of pro-angiogenic and -osteogenic factors. Notably, while we demonstrated the mTORC2-SGK1 signaling axis as a critical downstream pathway for ERK during late stage differentiation, the ERK MAPK pathway is likely to have additional substrates that contribute to its regulation of osteogenesis. Since WNT ligands can activate both the ERK and mTOR pathways independently to regulate skeletal development and homeostasis (Chen & Long, 2018, Karner & Long, 2017, Shim, Greenblatt et al., 2013), the ERK and mTORC2-SGK1 signaling axis may also function in parallel at different stages of osteogenesis.

ERK activation is dynamically changed in response to various extracellular stimuli during tissue development, homeostasis, and regeneration (Lavoie, Gagnon et al., 2020, Patel & Shvartsman, 2018). The ERK MAPK pathway plays a critical role in the regulation of cell quiescence/proliferation, differentiation, and commitment of adult stem cells and progenitors, as ERK activation promotes cell proliferation while suppressing cell differentiation (Krishnan, Kumar et al., 2021). However, very few *in vivo* studies have reported dynamics of ERK signaling in the skeletal system. This study demonstrates distinct roles of the ERK MAPK pathway at different stages of osteogenesis. In particular, during late-stage osteogenic differentiation, ERK activation is important for mitochondria-mediated glutamine metabolism rather than glucose metabolism (Fig 4). Previous studies have shown that osteogenic precursors not only consume glutamine to generate ATP but also synthesize metabolites to promote osteoblast viability and matrix mineralization (Yu, Newman et al., 2019). Our findings reveal that impaired ERK signaling enhances mitochondria-mediated glutamine metabolism, thereby promoting osteogenic differentiation. Remarkably, this can be reversed by the mTOR inhibitor rapamycin *in vitro* and *in vivo*. Since systemic administration of rapamycin increases bone anabolic activity in both normal bone mass and osteopenic conditions (Bateman, Sampurno et al., 2019, Chagin, 2016, Izawa, Martin et al., 2017), rapamycin-induced attenuation of osteogenic differentiation and bone anabolic activity in *Mek1/2*-deficient mice and osteoblasts is unlikely to result from non-specific or general effects of rapamycin unrelated to the specific defects in *Mek1/2*-deficient mice. Together with previous studies showing the association of the mTOR pathway with energy metabolism during skeletal development and homeostasis (Chen & Long, 2018), our data suggest that the ERK MAPK pathway controls energy metabolism in committed osteoblasts via the mTOR pathway.

This study also justifies an examination of the clinical impact of ERK pathway inhibitors on bone mass and fracture risk, as the biphasic complex contribution of the ERK pathway to bone formation makes empiric study of this question important. This is especially true given that the ERK pathway is the most frequently mutated pathway in cancer (Guo, Pan et al., 2020), and thus ERK inhibitors are important tools for targeted therapy in a wide range of tumor types. This study also shows that dual treatment with ERK and mTOR inhibitors may be associated with a strong decrease in bone mass, as mTOR inhibition would ablate the anabolic effect of ERK inhibition in late-stage osteoblasts, leaving only the negative effects of ERK on early osteogenic differentiation and commitment. In mice, trametinib treatment increased bone mass in osteoporotic bone (Fig 1A-C), indicating that the pro-anabolic effect of ERK inhibition in mature osteoblasts is dominant over the anti-anabolic effects on early-stage osteoblasts. However, the relative dominance of these opposing early versus late-stage effects may depend highly on dose and duration of therapy, in addition to clinical context and patient demographics. Thus, empirical study of the clinical effects of trametinib and other ERK pathway inhibitors will be important, especially given that many cancer patients being treated with these agents will be at increased risk for skeletal fractures due to the effects of either cancer, chemotherapy, or patient demographics impacting bone mass. Given the emerging role of osteoblasts in promoting and maintaining skeletal metastases, our finding that the ERK pathway has complex effects on early- and late-stage osteoblast lineage cells also suggests that further examination of the impact of ERK pathway inhibition of skeletal metastases will be important to understand the risks and benefits of ERK pathway inhibition (Bado, Zhang et al., 2021, Swami, Johnson et al., 2017, Zhang, Bado et al., 2021).

## Materials and methods

### Cell culture, antibodies, and reagents

Human bone marrow-derived mesenchymal stromal cells were purchased from Cyagen Biosciences and were maintained in growth medium (HUXMX- 90011) and cultured in osteogenic medium (GUXMX-90021). Primary osteoblasts were isolated from the calvaria of 5-day-old mouse pups using collagenase type II (50 mg/ml, Worthington, LS004176)/dispase II (100 mg/ml, Roche, 10165859001). *Mek1/2*-deficint osteoblasts were obtained from calvaria of 5-day-old *Mek1^fl/fl^Mek2^-/-^* neonates, transduced with lentiviruses expressing control or cre-recombinase, and selected with puromycin. Osteoblasts were maintained in α-minimal essential medium (Gibco) containing 10% fetal bovine serum (FBS, Corning), 2 mM L-glutamine (Corning), 1% penicillin/streptomycin (Corning) and 1% nonessential amino acids (Corning) and differentiated with ascorbic acid (200 μM, Sigma, A8960) and β-glycerophosphate (10 mM, Sigma, G9422).

Antibodies specific to MEK1 (12671), P-ERK1/2 (4376), P-p70S6K (9205), P-4EBP1 (2855), P-AKT (4060), P-NDRG1 (5482), and RUNX2 (12556) were purchased from Cell signaling. Antibodies specific to P-SGK1 (44-1264G) and GAPDH (CB1001) were purchased from Thermo Scientific and EMD Millipore, respectively. Trametinib (S2673) and BPTES (5301) were purchased from Selleck Chemicals and Tocris Bioscience, respectively. Rapamycin (sc-3054) for cell culture was purchased from Santa Cruz Biotechnology. All constructs encoding shRNAs were purchased from Sigma.

### Mice

*Mek1^fl/fl^* mice and *Mek2^-/-^* mice were generated as previously reported, respectively (Belanger et al., 2003, Bissonauth et al., 2006) and maintained on a 129/SvEv background. To generate mature osteoblast/osteocyte-specific double knockout (*Mek1^Dmp1^Mek2^-/-^*), *Mek1^fl/fl^ Mek2^-/-^* mice were crossed with *Dmp1*-*Cre* (Lu et al., 2007). Sex and age-matched littermates were used as controls for all skeletal analyses. To generate mice that harbored inducible deletion of *Mek1* in mature osteoblasts, *Mek1^fl/fl^Mek2^-/-^* mice were crossed with osteocalcin-*CreERT* (Park, Spencer et al., 2012) (*Mek1^Ocn-Ert^Mek2^-/-^*). For postnatal activation of *CreERT*, 75 mg/kg tamoxifen (Sigma, T5648) in corn oil (Sigma) was injected intraperitoneally into 8-week-old mice once a day, for 5 consecutive days. Rapamycin (R-5000, LC laboratories) was dissolved in ethanol and diluted in filter-sterilized vehicle (5% Tween-80, 5% PEG-40, 0.9% NaCl) immediately prior to intraperitoneal injection. Vehicle or 4 mg/kg rapamycin was administered every other day for 6 weeks. All animals were used under the NIH Guide for the Care and Use of Laboratory Animals and were handled according to the animal protocol approved by the University of Massachusetts Medical School Institutional Animal Care and Use Committee (IACUC).

### Ovariectomy (OVX)-induced bone loss

To induce postmenopausal osteoporosis, 12-week-old female mice were anesthetized and bilaterally ovariectomized or subjected to a sham operation. All mice were then randomly assigned to treatment with vehicle or trametinib (0.6 mg/kg) (Hu-Lieskovan, Mok et al., 2015). Inhibitor or vehicle treatment was given daily by oral gavage. After 8 weeks, mice were euthanized and subjected to bone analysis.

### Bone mechanical testing

To determine bone mechanical properties, 8-week-old male *Mek2^-/-^* (*Ctrl*) and *Mek1^Dmp1^Mek2^-/-^* (*dKO^Dmp1^*) femora were tested, as previously described (Melville, Kelly et al., 2014). Right femora were loaded to failure in three-point bending in the anterior-posterior direction with a span length of 6 mm at a rate of 0.1 mm/s (858 Mini Bionix: MTS, Eden Prairie, MN, USA). Bending strength was calculated based on the force and displacement data from the tests, and mid-shaft geometry was measured with microCT.

### MicroCT

MicroCT analysis was performed as previously described (Kim, Yang et al., 2020). Sex-matched and control littermates were used, and analysis was performed by an investigator blinded to the genotypes of the animals under analysis. Femurs excised from the indicated mice were scanned using a microCT 35 (Scanco Medical) with a spatial resolution of 7 μm. For trabecular bone analysis of the distal femur, an upper 2.1 mm region beginning 280 μm proximal to the growth plate was contoured. For cortical bone analysis of femur and tibia, a midshaft region of 0.6 mm in length was used. MicroCT scans of skulls and lumbar vertebrae (L4) were performed using isotropic voxel sizes of 12 μm. 3D reconstruction images were obtained from contoured 2D images by methods based on distance transformation of the binarized images. All images presented are representative of the respective genotypes (n>5).

### Histology, dynamic histomorphometry, and immunofluorescence

For histological analysis, hindlimbs were dissected from the mice, fixed in 10% neutral buffered formalin for 2 days, and decalcified by daily changes of 0.5 M tetrasodium EDTA (pH 7.4) for 3 to 4 weeks. Tissues were dehydrated by passage through an ethanol series, cleared twice in xylene, embedded in paraffin, and sectioned at 7 μm thickness along the coronal plate from anterior to posterior. Decalcified femoral sections were stained with hematoxylin and eosin (H&E) or tartrate-resistant acid phosphatase (TRAP). To detect apoptotic osteocytes in bone tissue, terminal deoxynucleotidyl transferase (TdT)-mediated dUTP-digoxigenin nick-end labeling (TUNEL) staining was performed using TUNEL assay kit (Abcam, ab206386) according to the manufacturer’s instructions.

For dynamic histomorphometry analysis, 25 mg/kg calcein (Sigma, C0875) and 50 mg/kg alizarin-3-methyliminodiacetic acid (Sigma, A3882) dissolved in 2% sodium bicarbonate solution were subcutaneously injected into mice at 5 day-intervals. After 2 day-fixation in 10% neutral buffered formalin, undecalcified femur samples were embedded in methylmethacrylate (Fukuda, Takeda et al., 2013). A region of interest was defined and bone formation rate/bone surface (BFR/BS), mineral apposition rate (MAR), osteoblast surface/bone surface (Ob.S/BS), osteoclast number/bone parameter (N.Oc/B.Pm), osteoclast surface/bone surface (Oc.S/BS), erosion surface/bone surface (ES/BS), osteocyte number/bone area (N.Ot/B.Ar), and empty lacunae were quantitated using a semiautomatic analysis system (OsteoMetrics, Atlanta, GA, USA). Measurements were taken on two sections/sample (separated by ∼25 μm) and summed prior to normalization to obtain a single measure/sample in accordance with ASBMR standards (Dempster, Compston et al., 2013, Parfitt, Drezner et al., 1987). This methodology underwent extensive quality control and validation and the results were assessed by a research specialist in a blinded fashion.

For immunofluorescence, decalcified and cryo-sectioned samples were incubated with antibodies for CD31 (1:100, BD Pharmingen, 553370) and endomucin (1:100; Santa Cruz, sc- 65495) were used as primary antibodies and Alexa Fluor 488 (1:400, Thermo, A21206) and Alexa Fluor 594 (1:400, Thermo, A11032) were used as secondary antibodies according to the manufacturer’s instructions.

### CTx-I measurement

Serum level of CTx-I was measured according to the manufacturer’s instructions (Immunodiagnostic Systems, AC-06F1).

### Transmission electron microscopy (TEM)

TEM images were obtained from JEOL JEM 1400. For TEM analysis of femoral bone, whole femur was fixed with 4% paraformaldehyde in 0.08 M Sorenson’s phosphate buffer at 4℃ overnight and decalcified with 10% EDTA (pH 7.2) for 10 days. Decalcified bones were embedded in epoxy resin after dehydration through an ethanol series and transferred to a transitional solvent, propylene oxide. Ultrathin sections of metaphysis area were examined for osteocyte analysis (Cheville & Stasko, 2014).

### Transcriptome analysis

Eight RNA-seq samples (two GM control, two GM *dKO*, two OIM control, and two OIM *dKO* samples) were mapped to the mouse reference genome (Mus_musculus.GRCm38.80) with STAR aligner (v.2.6.1b) (Dobin, Davis et al., 2013, Dobin & Gingeras, 2015). After mapping, read counts were generated by using HTSeq-count (v.0.11.3) (Anders, Pyl et al., 2015). The read counts were used for a differential expression analysis between control and *dKO* groups using DESeq2 (v.1.28.1) (Love, Huber et al., 2014) with the ashr shrinkage estimator (v.2.2.47) (Stephens, 2017). Statistically significantly expressed genes were determined as having absolute log-fold change larger than 1.5 and having a P-value less than 0.005. For upregulated genes (log fold change (LFC)>0; n=152 for GM and n=2048 for OIM) and downregulated genes (LFC<0; n=415 for GM and n=1576 for OIM) in *dKO* samples, gene ontology analysis was performed with enrichGO function in clusterProfiler package (v.3.18.1) (Yu, Wang et al., 2012) with the genome-wide annotation for mouse (org.Mm.eg.db; v.3.12) in Bioconductor.

### Analysis of cell proliferation

Cell proliferation was assessed by alamarBlue^TM^ staining (Thermo Scientific, DAL1100) or Bromodeoxyuridine (BrdU) incorporation assay (Abcam, ab126556).

### Collection of conditioned medium and osteoblast and endothelial cell functional assay

Conditioned medium (CM) was collected from control or *dKO* osteoblasts at day 6 of osteogenic culture. Briefly, cells were washed with PBS twice and incubated with serum free medium for 24 hours. Mouse endothelial progenitor outgrowth cells (EPOCs) were obtained from BioChain (7030031) and cultured in growth medium (BioChain Z7030035). For capillary tube formation assay, EPOCs (30000 cells/well) were seeded in a 96-well plate pre-coated with Matrigel (BD) and incubated with conditioned medium (CM) or FGF2. After 5 hours of incubation at 37 °C, the number of tube branches in each well was quantified by counting four random fields per well as previously described (Xu et al., 2018). Alternatively, CM was added to osteogenic culture of wildtype BMSCs and osteogenic differentiation was assessed after 6 days of culture.

### Measurement of glutamate and ATP production

Extracellular glutamate release was determined according to manufacturer’s protocols (Promega, J8021). Alternatively, intracellular ATP levels were determined by reaction of ATP with recombinant firefly luciferase and its substrate D-luciferin according to the manufacturer’s instructions (Thermo Scientific, A22066).

### Analysis of mitochondrial DNA copy number

The mitochondrial copy number was determined by mitochondrial DNA (mtDNA)/nuclear DNA (nDNA) using RT-PCR for mt-ND1/*Hk2* or mt- 16srRNA/*Hk2,* as previously reported (Quiros, Goyal et al., 2017). Sequences are provided in Table EV1.

### Assessment of mitochondrial number and membrane potential

Cellular mitochondria were stained with Mito Tracker (Biotium, 70075) and detected by confocal microscopy (Leica SP8). Alternatively, flow cytometery was used to assess mitochondrial number and membrane potential of the cells treated with Mito Tracker (Biotium, 70075) and TMRE (tetramethylrhodamine ethyl ester, Biotium, 70005), respectively, according to the manufacturers’ protocols. Flow cytometry analysis was performed on an LSR II (BD Biosciences).

### Measurement of oxygen consumption rate and intracellular ROS levels

Intracellular oxygen consumption rate (Cayman, 600800) and cellular reactive oxidative stress (ROS) levels (C10422, Invitrogen) were measured according to the manufacturers’ protocols.

### Luciferase reporter assay

RUNX2-responsive reporter gene (OG2-luc) and *Renilla* luciferase vector (Promega) were transfected into mouse calvarial osteoblasts using the Effectene transfection reagent (Qiagen). After 48 hours, dual luciferase assay was performed according to the manufacturer’s protocol (Promega) and OG2 luciferase activity was normalized to *Renilla*.

### RT-PCR and immunoblotting

To prepare bone RNA samples from mouse limbs, the hindlimbs were dissected and skin/muscle tissues were removed. The remaining tibias were chopped and homogenized. Total RNA from cells or tissues were extracted using QIAzol (QIAGEN) and cDNA was synthesized using the High-Capacity cDNA Reverse Transcription Kit from Applied Biosystems. Quantitative RT-PCR was performed using SYBR® Green PCR Master Mix (Bio-Rad, Hercules, CA) with Bio-Rad CFX Connect Real-Time PCR detection system. mRNA levels were normalized to the housekeeping gene Ribosomal protein, large, P0 (*Rplp0*). The primers used for PCR are described in Table EV1.

For immunoblotting, cell lysates were prepared in lysis buffer (50 mM Tris-HCl [pH 7.8], 150 mM NaCl, 1% Triton X-100, 1 mM dithiothreitol [DTT], 0.2% sarkosyl acid and protease inhibitor cocktail [Sigma, P8340]). Protein samples were subjected to SDS-PAGE and transferred to Immobilon-P membranes (Millipore). Membranes were immunoblotted with the indicated antibodies and developed with ECL (Thermo Scientific). Immunoblotting with antibody specific to GAPDH was used as a loading control.

### Statistics and reproducibility

All experiments were performed a minimum of two to three times. For histological staining, flow cytometry, and immunoblotting, representative images are shown. All data are shown as the mean ± standard deviation (SD). We first performed the Shapiro-Wilk normality test for checking normal distributions of the groups. For comparisons between two groups, a two-tailed unpaired Student’s t-test was used if normality tests passed and a Mann-Whitney test was used if normality tests failed. For the comparisons of three to six groups, we used one-way ANOVA if normality tests passed, followed by Tukey’s multiple comparison test for all pairs of groups. GraphPad PRISM software (ver.9.0.2, La Jolla, CA) was used for statistical analysis. P<0.05 was considered statistically significant.

### Data availability

Data supporting the findings of this manuscript are available from the corresponding author upon reasonable request.

### Author Contributions

J.M.K. designed, executed, and interpreted the experiments. Y.S.Y. and S.C. performed histology, immunohistochemistry, dynamic histomorphometry, and microCT analyses. J.H. and H.C. performed transcriptome analysis. M.M. performed bone mechanical tests. R.X. performed angiogenic assays. M.B.G. assisted with skeletal analysis of mice. J.H.S. supervised the research and participated in the manuscript preparation. All authors reviewed and approved the final version of the manuscript.

## Acknowledgments

We would like to thank Dr. Hwanhee Oh for help with the skeletal analysis. We also thank Oksun Lee and Jihea Kim for experimental support and the many individuals who provided valuable reagents. This project was supported by NIH-NIAMS R21AR077557 and AAVAA Therapeutics.

M.B.G. holds a Career Award for Medical Scientists from the Burroughs Wellcome Fund, NIH support under award R01AR075585, a Novartis Institutes for Biomedical Research Global Scholars Award, and a Pershing Square Sohn Cancer Research Alliance award.

## Competing Financial Interests

J.H.S. is a scientific co-founder of the AAVAA Therapeutics and holds equity in this company.

Other authors declare no competing interests.

