## Supplemental information for "Impaired ERK MAPK activation in mature osteoblasts enhances bone formation via the mTOR pathway"

Department of Medicine/Division of Rheumatology

#### Expanded View Table 1. Primer sequences

| Gene | Forward | Reverse |
|---|---|---|
| Human *RUNX2* | TTACTTACACCCCGCCAGTC | CACTCTGGCTTTGGGAAGAG |
| Human *SP7* | TTACAAGCACTAATGGGCTCCT | GTAGACACTGGGCAGACAGTCA |
| Human *RPLP0* | GGAATGTGGGCTTTGTGTTC | TGCCCCTGGAGATTTTAGTG |
| Mouse *Ibsp* | CAGGGAGGCAGTGACTCTTC | AGTGTGGAAAGTGTGGCGTT |
| Mouse *Bglap* | GCAGCACAGGTCCTAAATAG | GGGCAATAAGGTAGTGAACAG |
| Mouse *Col1a1* | ACTGTCCCAACCCCCAAAG | ACGTATTCTTCCGGGCAGAA |
| Mouse *Map2k1* (*Mek1*) | ACTGCCCAGTGGAGTATTCAGT | TGTACCATGAGCTGCTTCAGAT |
| Mouse *Map2k2* (*Mek2*) | CATCAGTGTAGGTCATGGGATG | AAACTCCTGGAAGTCTGAGCTG |
| Mouse *Dmp1* | GAAAGCTCTGAAGAGAGGACGG | CCTCTCCAGATTCACTGCTGTC |
| Mouse *Phex* | CTGGCTGTAAGGGAAGACTCCC | GCTCCTAAAAGCACAGCAGTGTC |
| Mouse *Sost* | CTTCAGGAATGATGCCACAGAGGT | ATCTTTGGCGTCATAGGGATGGTG |
| Mouse *Gls* | GGCAAAGGCATTCTATTGGA | CTTGGCTCCTTCCCAACATA |
| Mouse *Pgc1α* (*Ppargc1a*) | CCCACAACTCCTCCTCATAAAG | TTGGGTACCAGAACACTCACTG |
| Mouse *Drp1* | GGGGTAAATTTCTTCACACCAA | TCAGGGCTTACCCCCTTATTAT |
| Mouse *Mfn1* | ATAGAAGATGGCATGGAAGAA | GCTGGAAGTAGTGGCTTCAAGT |
| Mouse *Sgk1* | GGGTGCCAAGGATGACTTTA | TGGGTTAAATGGGGGTGTAA |
| Mouse *Rplp0* | TGGCCAATAAGGTGCCAGCTGCTG | CTTGTCTCCAGTCTTTATCAGCTGCAC |
| Mouse *Hk2* | GCCAGCCTCTCCTGATTTTAGTGT | GGGAACACAAAAGACCTCTTCTGG |
| mt-*ND1* | CTAGCAGAAACAAACCGGGC | CCGGCTGCGTATTCTACGTT |
| mt-*16srRNA* | CCGCAAGGGAAAGATGAAAGAC | TCGTTTGGTTTCGGGGTTTC |

#### Expanded View Figure Legends

##### Expanded View Figure 1. Osteogenic gene expression in human BMSCs

Human BMSCs were cultured under osteogenic conditions, and mRNA levels of *RUNX2* and *SP7* were analyzed by RT-PCR at day 0 and 8 in culture. A two-tailed unpaired Student's t-test for comparing two groups (error bars represent the SD of biological replicates).

##### Expanded View Figure 2. Analysis of skeletal phenotypes in $Mek1^{Dmp1}Mek2^{-/-}$ mice

(A) TRAP-stained longitudinal sections of 8-week-old *WT* and $dKO^{Dmp1}$ femurs. Representative images (top) and the relative quantification (bottom) are shown. N.Oc/B.pm, osteoclast number per bone perimeter; ES/BS, erosion surface per bone surface. Scale bar, 20 μm (top).

(B) Serum levels of cross-linked C-telopeptide of type 1 collagen (CTx-I) in 8-week-old *WT* and $dKO^{Dmp1}$ male mice were measured by ELISA.

(C) mRNA levels of *Dmp1*, *Phex*, and *Sost* in 8-week-old *WT* and $dKO^{Dmp1}$ tibial bone RNA were assessed by RT-PCR. Data are representative of two or three independent experiments (A [top], C) or pooled from two experiments (A [bottom], B). A two-tailed unpaired Student's t-test for comparing two groups (A-C; error bars represent the SD of biological replicates).

##### Expanded View Figure 3. Characterization of *Mek1/2*-deficient osteoblast-lineage cells

(A) Mouse $Mek1^{fl/fl}Mek2^{+/+}$ and $Mek1^{fl/fl}Mek2^{-/-}$ OBs were infected with control or CRE recombinase-expressing lentiviruses; $Mek1^{fl/fl}Mek2^{+/+}$ OBs with control (WT) or CRE (*Mek1 KO)*,

*Mek1* $^{fl/fl}$*Mek2*$^{-/-}$ OBs with control (*Mek2 KO)* or CRE *(Mek1/2 dKO)*. Puromycin-selected OBs were cultured under osteogenic conditions and *Col1a1* gene expression was determined at day 6. (B, C) *Ctrl* (*Mek2 KO*) and *dKO* (*Mek1/2 dKO)* OBs were cultured in the presence of growth medium (GM) or osteogenic induction medium (OIM) and ALP activity (B) and osteogenic gene expression (C) were assessed at day 6 in culture.

(D-G) Transcriptome analysis of *Ctrl* and *dKO* OBs 6 days after GM or OIM culture. Biological process output of gene ontology analysis was performed in both GM (D) and OIM (E) group for downregulated genes in *dKO* OBs relative to *Ctrl* OBs. The color indicates adjusted P-value as estimated by the Benjamini–Hochberg method with the threshold of significance $p = 0.05$ and $q = 0.005$. Volcano plots showing the gene expression for up/downregulated genes in *dKO* OBs relative to *Ctrl* OBs after GM (F) or OIM (G) culture. Dots indicate upregulated (red) and downregulated genes (blue). The top 10 up/downregulated genes were labeled. Data are representative of three independent experiments (A-C). An ordinary one-way ANOVA with Dunnett's multiple comparisons test (A) or a two-tailed unpaired Student's t-test for comparing two groups (B, C) (A-C; error bars represent the SD of biological replicates).

**Expanded View Figure 4. Effects of *Mek1/2* deletion on RUNX2 expression and transcriptional activity in osteoblasts**

(A) Immunoblotting analysis showing protein levels of RUNX2 in *Ctrl* and *dKO* OBs. GAPDH was used as a loading control.

(B) *Ctrl* and *dKO* OBs were transfected with the OG2-luc reporter gene along with *Renilla*. After 48 hours, luciferase activity was measured and normalized to *Renilla* activity. Data are representative of three independent experiments. A two-tailed unpaired Student's t-test was used to compare groups (B; error bars represent the SD of biological replicates).

**Expanded View Figure 5. ERK inhibition enhances mitochondrial function in osteoblasts**

(A) Mouse wildtype OBs were treated with vehicle or trametinib (0.5 μM) under osteogenic conditions and 6 days later, mitochondrial DNA (mtDNA) copy number was determined by mitochondrial DNA (mtDNA, mt-ND1 or mt-16sRNA) to nuclear DNA (nDNA, *Hk2*) ratio using RT-PCR.

(B) Mitochondrial mass (left) and membrane potential (right) in vehicle- or 0.5 μM trametinib-treated OBs were analyzed using Mitotracker and TMRE staining, respectively, at day 4 of the osteogenic culture. Numbers indicate median fluorescence intensity. Data are representative of three independent experiments. A two-tailed unpaired Student's t-test for comparing two groups (A; error bars represent the SD of biological replicates).

**Expanded View Figure 6. Rapamycin inhibits both mTORC1 and mTORC2 pathways**

Immunoblotting analysis showing protein levels of P-AKT, P-SGK1 and P-p70S6K in *dKO* OBs. GAPDH was used as a loading control. Data are representative of three independent experiments.

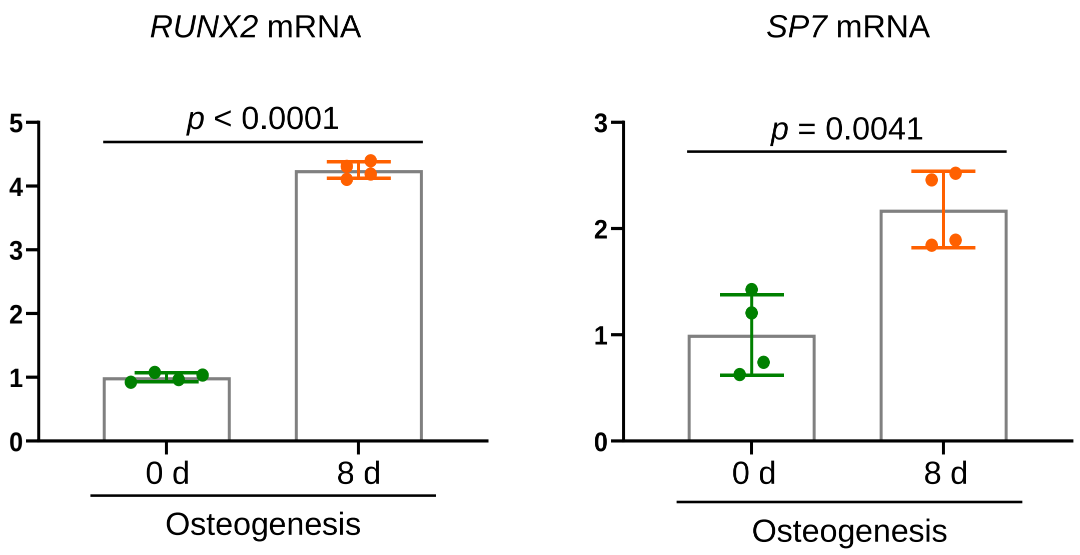

*RUNX2* mRNA
*p* < 0.0001
5
4
3
2
1
0
0 d
8 d
Osteogenesis
*SP7* mRNA
*p* = 0.0041
3
2
1
0
0 d
8 d
Osteogenesis


**Figure EV1**

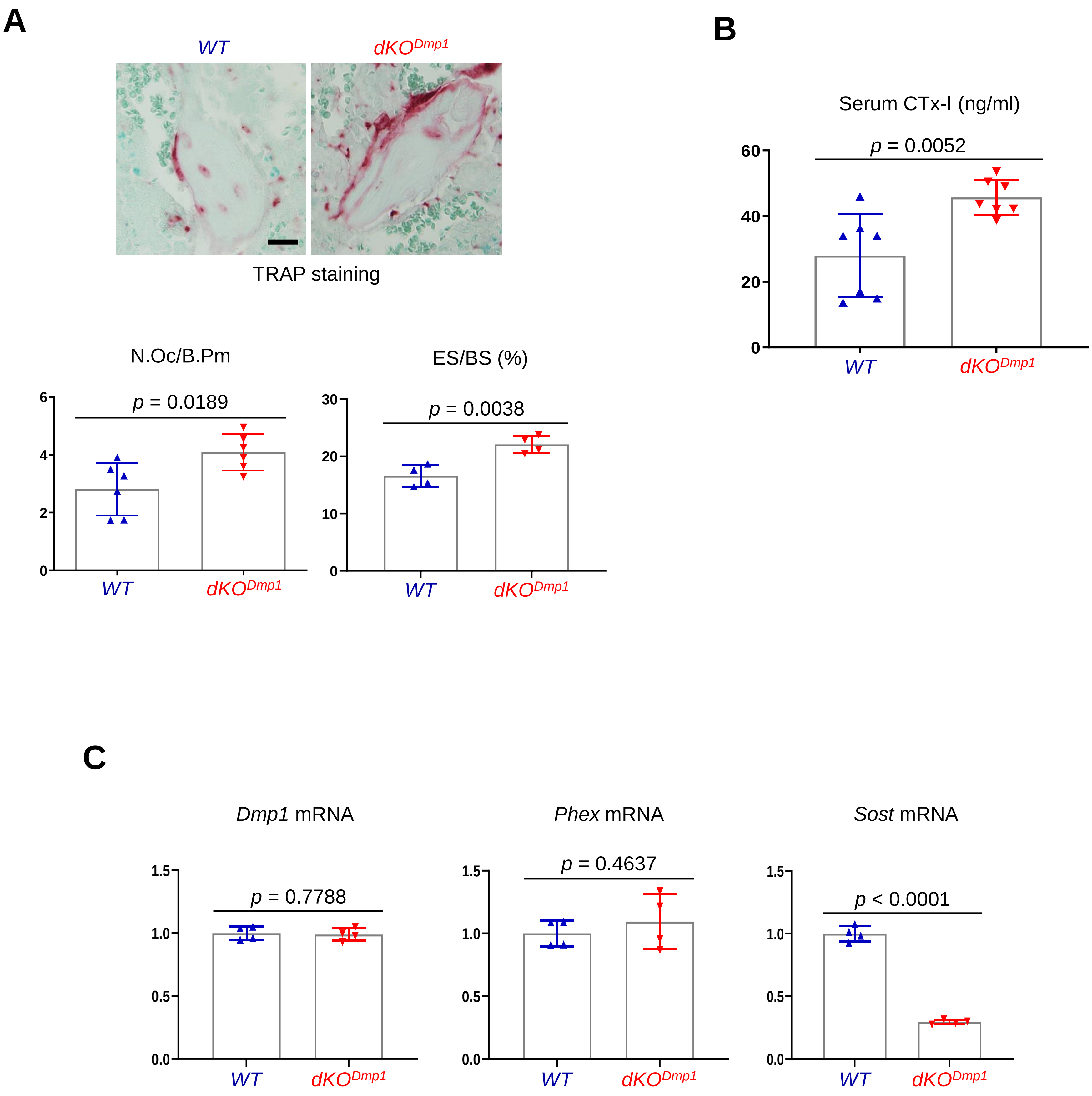
A
WT
dKO^Dmp1
TRAP staining
N.Oc/B.Pm
p = 0.0189
ES/BS (%)
p = 0.0038
B
Serum CTx-I (ng/ml)
p = 0.0052
C
Dmp1 mRNA
p = 0.7788
Phex mRNA
p = 0.4637
Sost mRNA
p < 0.0001

**Figure EV2**

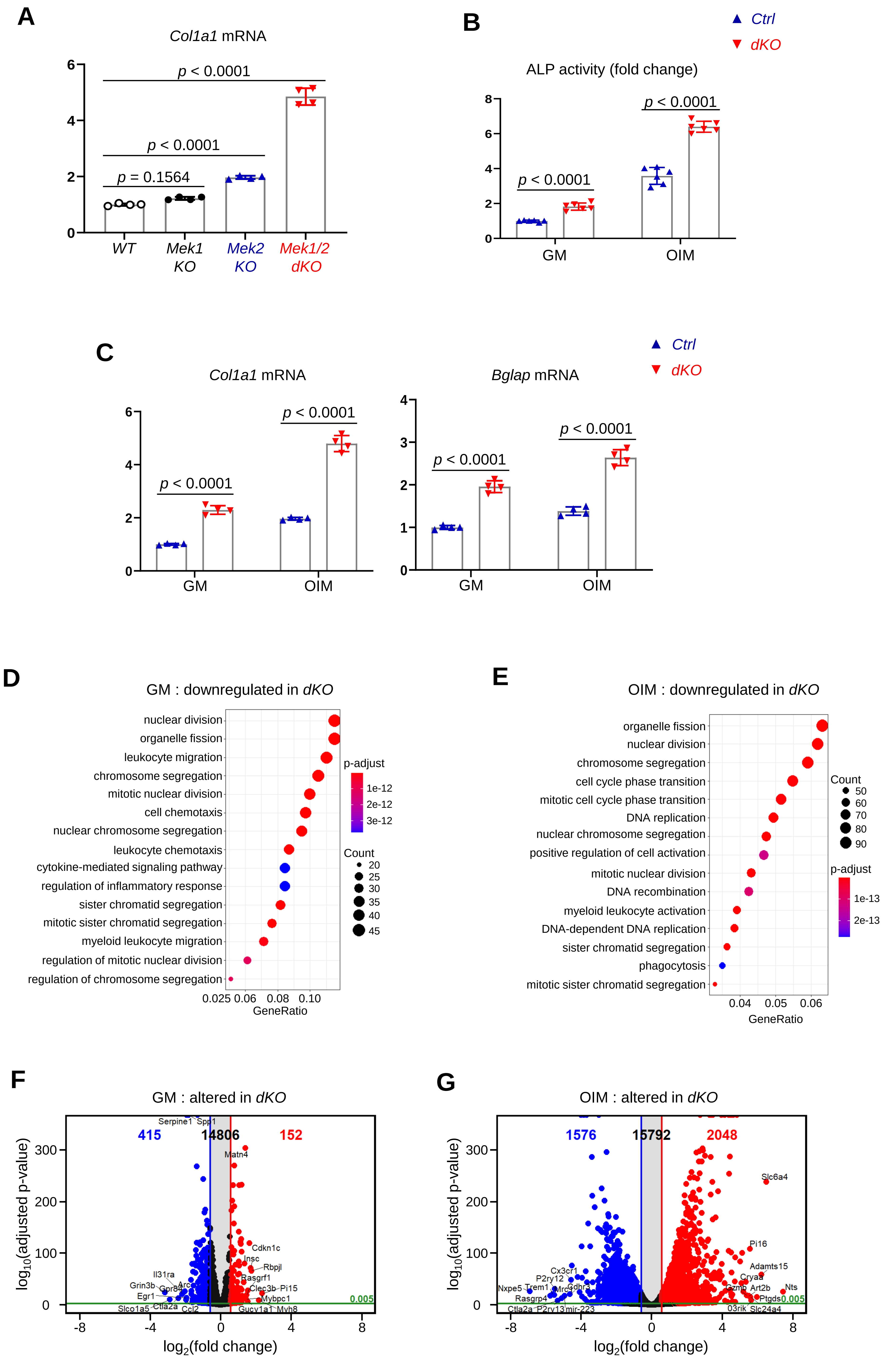

A
Col1a1 mRNA
p < 0.0001
p < 0.0001
p = 0.1564
WT
Mek1 KO
Mek2 KO
Mek1/2 dKO
B
Ctrl
dKO
ALP activity (fold change)
p < 0.0001
p < 0.0001
GM
OIM
C
Col1a1 mRNA
Bglap mRNA
p < 0.0001
GM
OIM
D
GM : downregulated in dKO
nuclear division
organelle fission
leukocyte migration
chromosome segregation
mitotic nuclear division
cell chemotaxis
nuclear chromosome segregation
leukocyte chemotaxis
cytokine-mediated signaling pathway
regulation of inflammatory response
sister chromatid segregation
mitotic sister chromatid segregation
myeloid leukocyte migration
regulation of mitotic nuclear division
regulation of chromosome segregation
p-adjust
Count
GeneRatio
E
OIM : downregulated in dKO
organelle fission
nuclear division
chromosome segregation
cell cycle phase transition
mitotic cell cycle phase transition
DNA replication
nuclear chromosome segregation
positive regulation of cell activation
mitotic nuclear division
DNA recombination
myeloid leukocyte activation
DNA-dependent DNA replication
sister chromatid segregation
phagocytosis
mitotic sister chromatid segregation
Count
p-adjust
GeneRatio
F
GM : altered in dKO
415
14806
152
log10(adjusted p-value)
log2(fold change)
G
OIM : altered in dKO
1576
15792
2048
log10(adjusted p-value)
log2(fold change)


**Figure EV3**

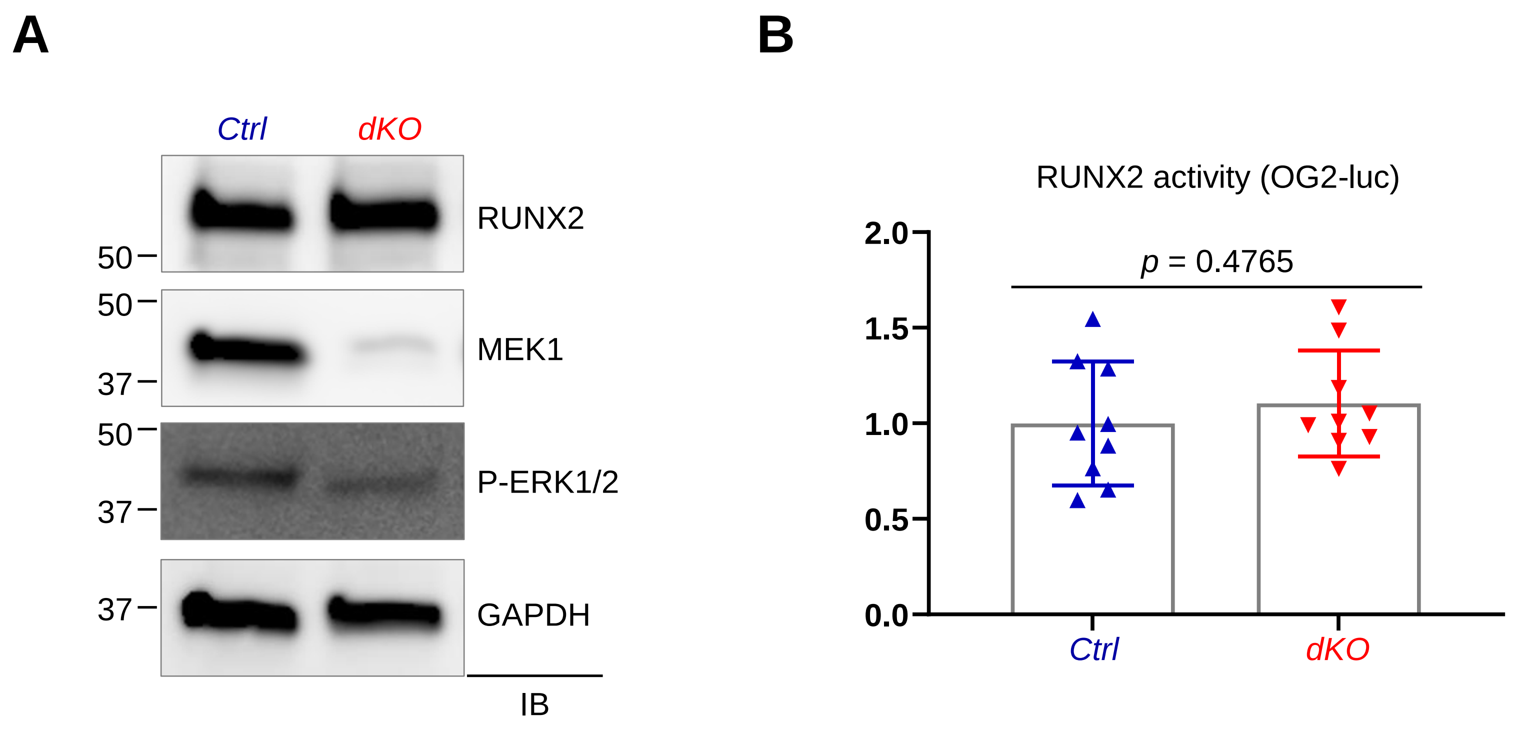

A
B
Ctrl
dKO
RUNX2
MEK1
P-ERK1/2
GAPDH
IB
50
50
37
50
37
37
RUNX2 activity (OG2-luc)
p = 0.4765
2.0
1.5
1.0
0.5
0.0
Ctrl
dKO


### Figure EV4

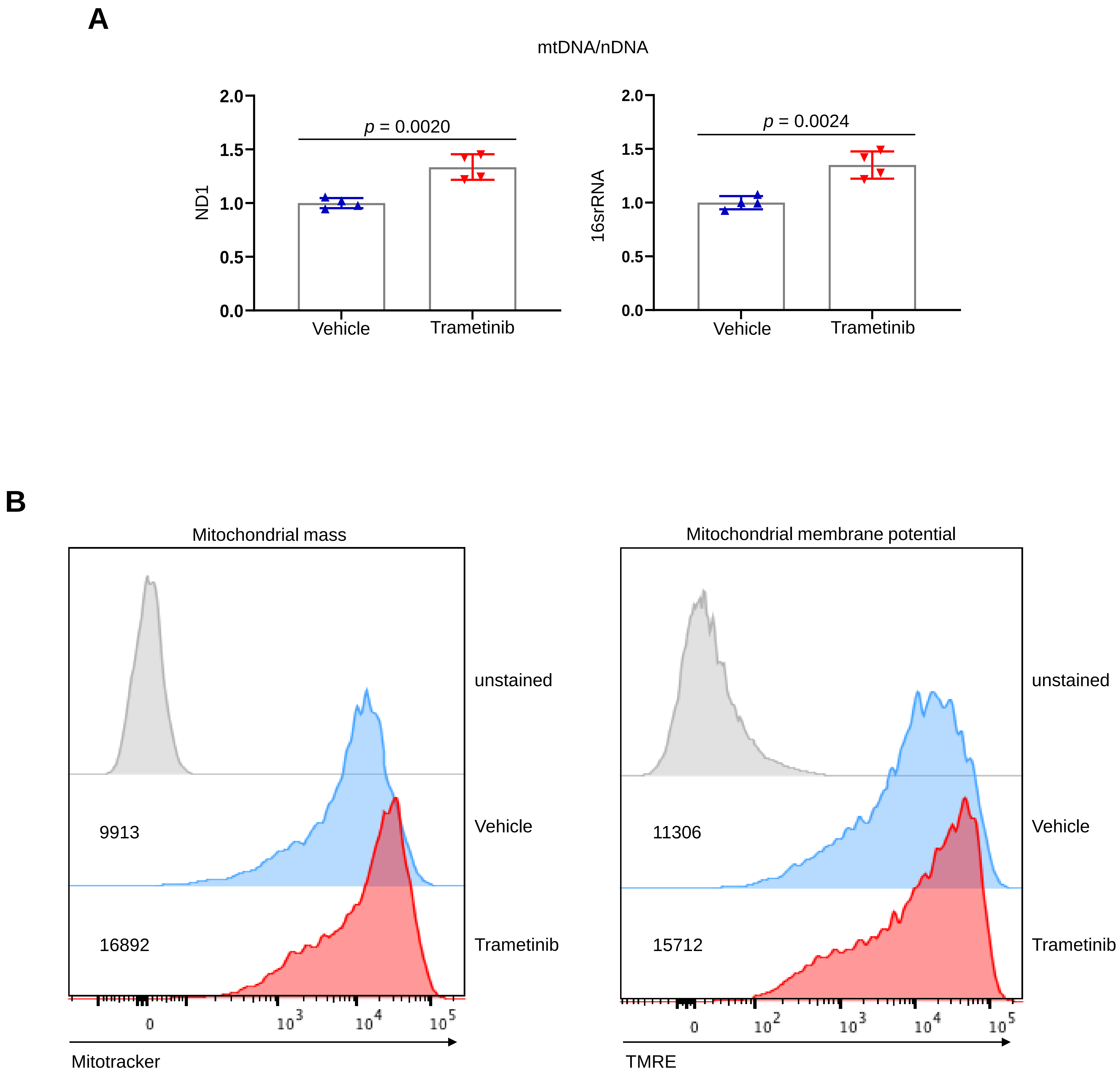
A
mtDNA/nDNA
p = 0.0020
ND1
Vehicle
Trametinib
p = 0.0024
16srRNA
Vehicle
Trametinib
B
Mitochondrial mass
unstained
Vehicle
Trametinib
9913
16892
Mitotracker
Mitochondrial membrane potential
unstained
Vehicle
Trametinib
11306
15712
TMRE

Figure EV5

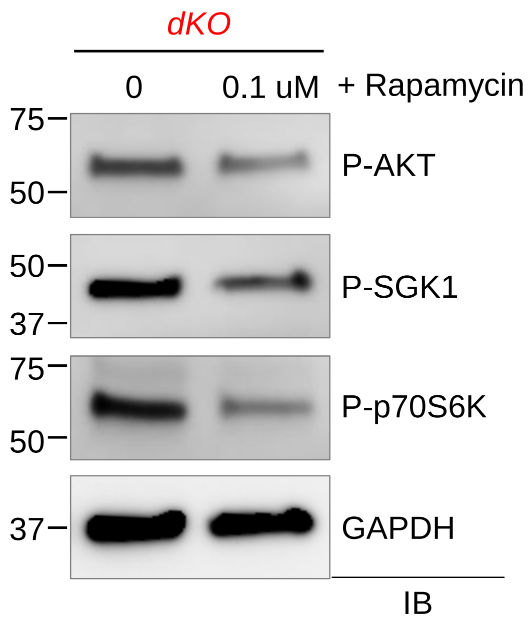

dKO
0
0.1 uM
+ Rapamycin
75
50
P-AKT
50
37
P-SGK1
75
50
P-p70S6K
37
GAPDH
IB


**Figure EV6**
